# A global map of root biomass across the world’s forests

**DOI:** 10.1101/2020.01.14.906883

**Authors:** Yuanyuan Huang, Phillipe Ciais, Maurizio Santoro, David Makowski, Jerome Chave, Dmitry Schepaschenko, Rose Abramoff, Daniel S. Goll, Hui Yang, Ye Chen, Wei Wei, Shilong Piao

## Abstract

Root plays a key role in plant growth and functioning. Here we combine 10307 field measurements of forest root biomass worldwide with global observations of forest structure, climatic conditions, topography, land management and soil characteristics to derive a spatially-explicit global high-resolution (~ 1km) root biomass dataset, including fine and coarse roots. In total, 142 ± 32 Pg of live dry matter biomass is stored below-ground, that is a global average root:shoot biomass ratio of 0.25 ± 0.10. Our estimations of total root biomass in tropical, temperate and boreal forests are 44-226% smaller than earlier studies^1–3^. The smaller estimation is attributable to the updated forest area, spatially explicit above-ground biomass density used to predict the patterns of root biomass, new root measurements and upscaling methodology. We show specifically that the root shoot allometry is one underlying driver that leads to methodological overestimation of root biomass in previous estimations.

---

Roots mediate nutrient and water uptake by plants, below-ground organic carbon decomposition, the flow of carbohydrates to mycorrhizae, species competition, soil stabilization and plant resistance to windfall^4^. The global distribution of root biomass is related to how much photosynthates plants must invest below-ground to obtain water, nitrogen and phosphorus for sustaining photosynthesis, leaf area and growth. Root biomass and activity also control the land surface energy budget through plant transpiration^4,5^. While Earth Observation data combined with field data enables the derivation of spatially explicit estimates of above-ground biomass with a spatial resolution of up to 30 meters over the whole globe^6,7^, the global carbon stock and spatial details of the distribution of below-ground root biomass (fine + coarse) relied on punctual measurements and coarse extrapolation so far, therefore remaining highly uncertain

More than twenty years ago, Jackson et al, 1996, 1997 ^1,2^ provided estimates of the average biomass density (weight per unit area) and vertical distribution of roots for 10 terrestrial biomes. Multiplying their average root biomass density with the area of each biome gives a global root biomass pool of 292 Pg, with forests accounting for ~68% of it. Saugier, et al. (2001) estimated global root biomass to be 320 Pg by multiplying biome-average root to shoot ratios (*R:S*) by shoot biomass density and the land area of each biome. Mokany, et al. (2006) argued that the use of mean *R:S* values at biome scale is a source of error because root biomass measurements are performed at small scales with roots having a high spatial heterogeneity and their size distribution spanning across several orders of magnitude, the fine roots being particularly difficult to sample^8,9^. With updated *R:S* and broader vegetation classes, they gave a higher global root biomass of 482 Pg. Robinson (2007) further suggested a 60% underestimation of *R:S*, which translated into an even higher global root biomass of 540-560 Pg. These studies provided a first order estimation of the root biomass for different biomes, but not of its spatial details and it is worth noting that numbers have increased with time.

An alternative approach to estimate root biomass is through allometric scaling, dating back to West, Brown and Enquist (1997, 1999)^6 7^ and Enquist and Niklas (2002). The allometric scaling theory assumes that biological attributes scale with body mass, and in the case of roots, an allometric equation verified by data takes the form of *R* ∝ *S*^*β*^ where *R* is the root mass, *S* the shoot mass and *β* a scaling exponent. Differently than in the studies listed above assuming the R:S ratio to be uniform, this equation implies that the *R:S* ratio varies with shoot size as β is not equal to one ^10–15^. Allometric equations also predict that smaller trees generally have a larger *R:S* with *β* < 1, which is well verified by measurement of trees of different sizes ^12–15^. The allometric equation approach was applied for various forest types, and the scaling exponent *β* was observed to differ across sites^16^, species^17^, age^13^, leaf characteristics^18^, elevation^19^, management status^20^, climatic conditions, such as temperature^21^, soil moisture and climatic water deficit ^20^, as well as soil nutrient content and texture^14^. Despite successful application of allometric equations for site- and species-specific studies^16^, their use to predict large-scale and global root biomass patterns appears to be challenging.

Here we use a new approach to upscale root biomass of trees at global scale based on machine learning algorithms trained by a large dataset of measurements and using as predictors high-resolution maps of tree density, above-ground biomass, soils and environmental drivers (Supplementary Tables 1, 2). Firstly, we collected 10307 *in-situ* measurements ^14,30,31^of the biomass of roots and shoots for individual woody plants (see Methods, Supplementary Data), covering 465 species across 10 biomes defined by The Nature Conservancy^22^ (Supplementary Figure 2). In biomes like savannas where trees and woody plants can be sparse, we estimate root biomass as the average for the woody plants present in that biome given a canopy cover threshold of 15% at a 30 m resolution globally^23^. In the root below-ground biomass estimates (BGB) we count both coarse and fine roots. We acknowledge the importance of understanding large scale temporal dynamics of fine root. As a first step, this study aims at the spatial pattern of total root biomass. We upscaled root biomass from individual plant level measurements rather than from stand-level data because a large number of primary data are collected for individual woody plants, and this approach allows us to account in both the training of machine learning models and their upscaling results, for the fact that root biomass depends on tree size or above-ground biomass^14,15,20^. We searched through a pool of 47 predicting variables that include above-ground biomass and other vegetation variables, edaphic, topographic, anthropogenic and climatic conditions (Supplementary Table 1). Different machine learning models were tested, and we selected the model that performs best on cross validation samples (see Methods for model selection criterion). The best model is a random forest (RF, see Methods) and we mapped global root biomass at a 1 km resolution through this model relying on 14 predicting gridded variables, including the shoot biomass of an average tree derived from shoot density (weight per area) ^24^ and tree density (number of trees per area)^25^, tree height^26^, soil nitrogen^27^, pH^27^, bulk density^27^, clay content^27^, sand content^27^, base saturation^27^, cation exchange capacity^27^, water vapor pressure^28^, mean annual precipitation^28^, mean annual temperature^28^, aridity^29^ and water table depth^30^ (see Supplementary Table 1 for detailed information and references). To estimate root biomass pools at global and biome scales, the mean root biomass of trees in each 1 km pixel was multiplied by a tree density map available at the resolution of 1 km from ref. ^25^ (see Methods)

## Results

We estimated a global total root biomass of 142±32 Pg (see Method for uncertainty estimation and Supplementary Figures 3, 4) when forest is defined as all areas with tree cover larger than 15% from the Hansen et al. (2013) tree cover map. The corresponding global weighted mean *R:S* is 0.25 ± 0.10. The root biomass spatial distribution generally follows the pattern of shoot biomass, but there are significant local and regional deviations as shown by Figure 1. 51% of the global tree root biomass comes from tropical moist forest, 14% from boreal forest, 12% from temperate broadleaf forest and 10% from woody plants in tropical and subtropical grasslands, savanna and shrublands (Supplementary Table 3). Given our use of a tree cover threshold of 15% at 30m resolution, our estimate ignores the roots of isolated woody plants present in arid or cold regions ^31^, as well as heterogeneous (e.g. urban or agriculture) landscapes and is possibly an under-estimate. Total root biomass decreases from 151 to 134 Pg when the canopy cover threshold used to define forest land is increased from 0% to 30%. The root biomass density per unit of forest area is highest in tropical moist forest, followed by temperate coniferous and Mediterranean forest (Figure 1, Supplementary Table 3). Cross validation showed a good match between predictions from our RF model and *in-situ* observations (Figures 2e, all data; Supplementary Figure 6, for each biome), with a coefficient of determination *R*^2^ of 0.85 and a median *R:S* similar to validation samples (0.35 from *in-situ* observation vs. 0.38 from prediction). Root biomass of tropical, temperate and boreal forests together is 44-226% lower compared to earlier studies (Table 1, Supplementary Table 5, see Supplementary Information Comparison with published results).

**Table 1.**
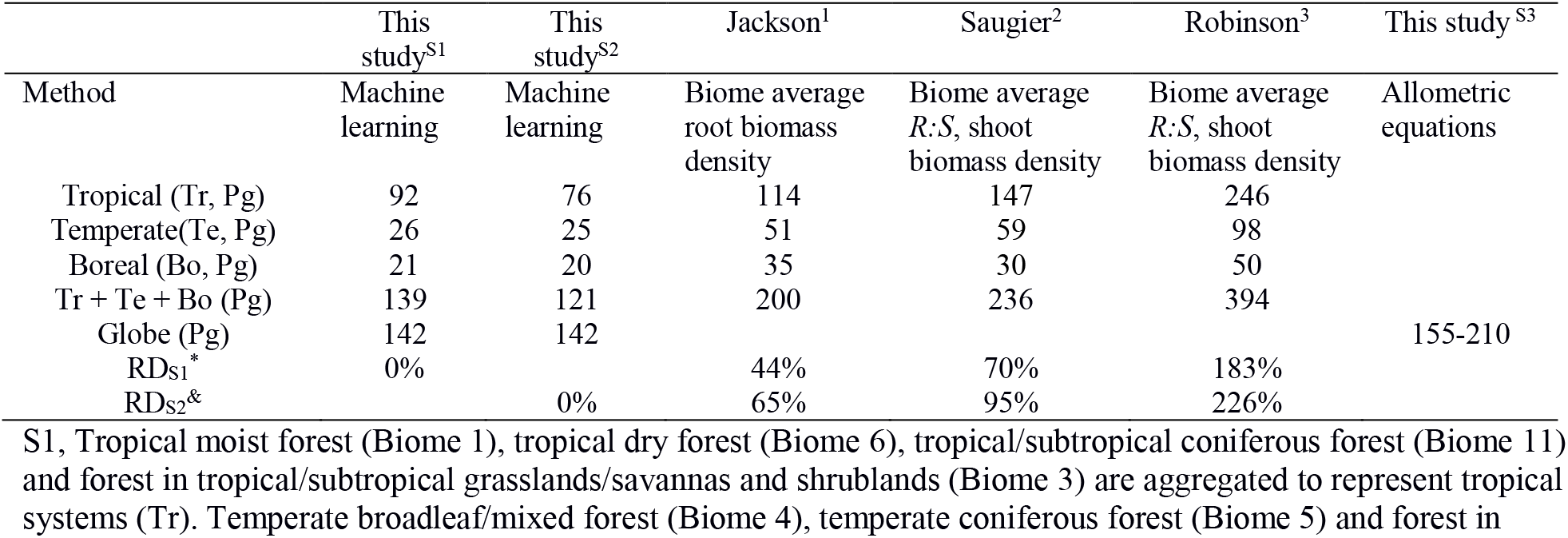
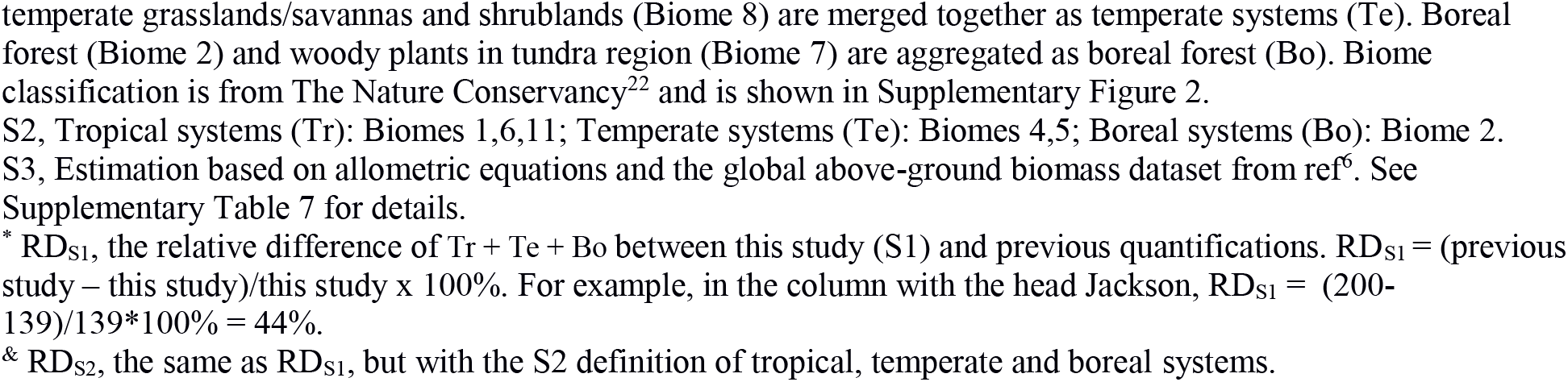
Comparison between studies quantifying root biomass in tropical, temperate and boreal forests.

|  | This study<sup>S1</sup> | This study<sup>S2</sup> | Jackson<sup>1</sup> | Saugier<sup>2</sup> | Robinson<sup>3</sup> | This study<sup>S3</sup> |
| --- | --- | --- | --- | --- | --- | --- |
| Method | Machine learning | Machine learning | Biome average root biomass density | Biome average $R:S$, shoot biomass density | Biome average $R:S$, shoot biomass density | Allometric equations |
| Tropical (Tr, Pg) | 92 | 76 | 114 | 147 | 246 |  |
| Temperate (Te, Pg) | 26 | 25 | 51 | 59 | 98 |  |
| Boreal (Bo, Pg) | 21 | 20 | 35 | 30 | 50 |  |
| Tr + Te + Bo (Pg) | 139 | 121 | 200 | 236 | 394 |  |
| Globe (Pg) | 142 | 142 |  |  |  | 155-210 |
| RDs<sup>S1</sup>* | 0% |  | 44% | 70% | 183% |  |
| RDs<sup>S2</sup>& |  | 0% | 65% | 95% | 226% |  |

S1, Tropical moist forest (Biome 1), tropical dry forest (Biome 6), tropical/subtropical coniferous forest (Biome 11) and forest in tropical/subtropical grasslands/savannas and shrublands (Biome 3) are aggregated to represent tropical systems (Tr). Temperate broadleaf/mixed forest (Biome 4), temperate coniferous forest (Biome 5) and forest in
S2, Tropical systems (Tr): Biomes 1,6,11; Temperate systems (Te): Biomes 4,5; Boreal systems (Bo): Biome 2.
S3, Estimation based on allometric equations and the global above-ground biomass dataset from ref<sup>6</sup>. See Supplementary Table 7 for details.
\* $RD_{S1}$ , the relative difference of Tr + Te + Bo between this study (S1) and previous quantifications. $RD_{S1} = (\text{previous study} - \text{this study}) / \text{this study} \times 100\%$ . For example, in the column with the head Jackson, $RD_{S1} = (200 - 139) / 139 \times 100\% = 44\%$ .
& $RD_{S2}$ , the same as $RD_{S1}$ , but with the S2 definition of tropical, temperate and boreal systems.

**Figure 1.**
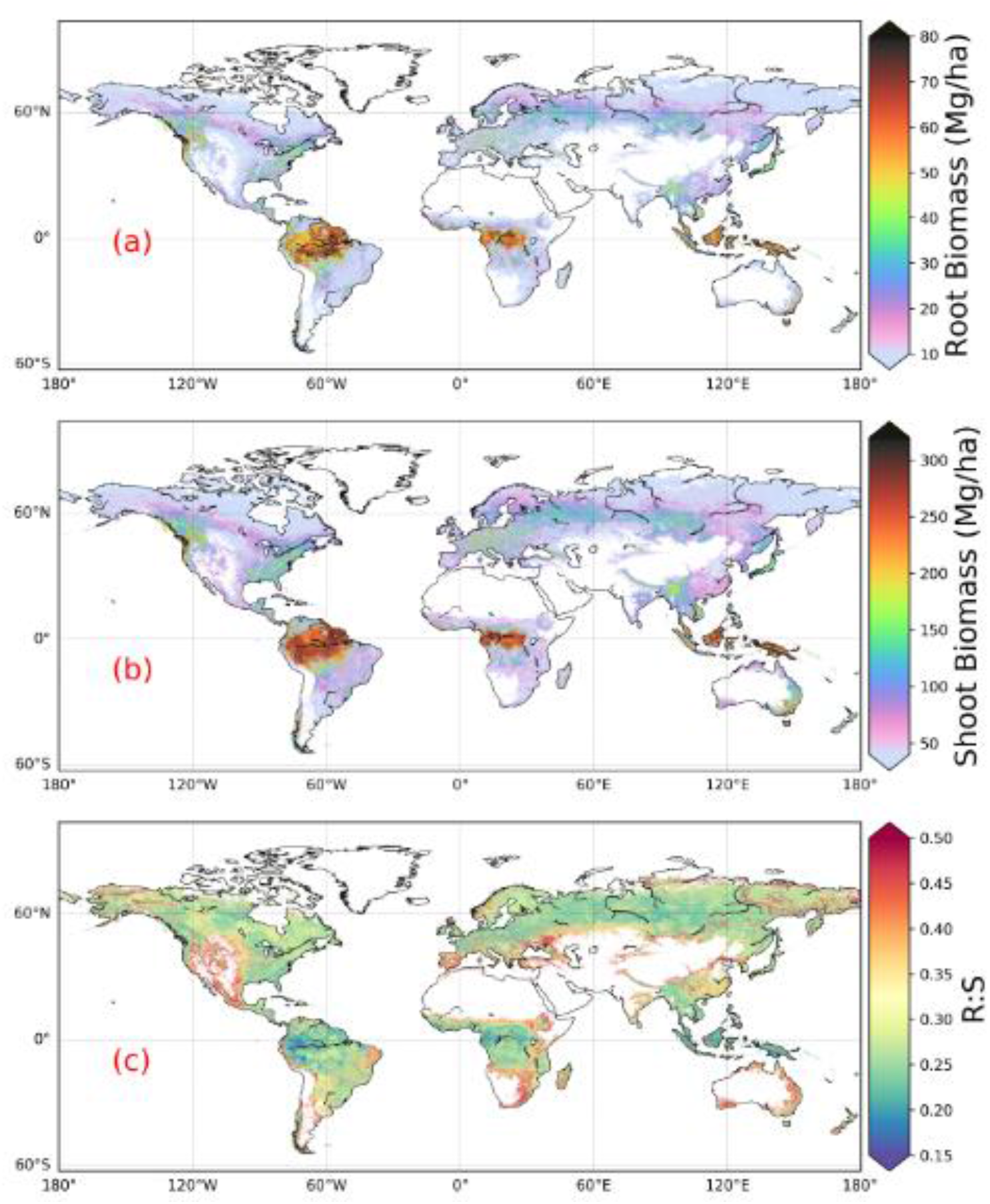
Global maps of forest root biomass generated through the random forest model (a), shoot biomass from GlobBiomass-AGB^6^ (b) and *R:S* (c). Forest is defined as an area with canopy cover > 15% from the Hansen et al. (2013) tree cover map.

**Figure 2.**
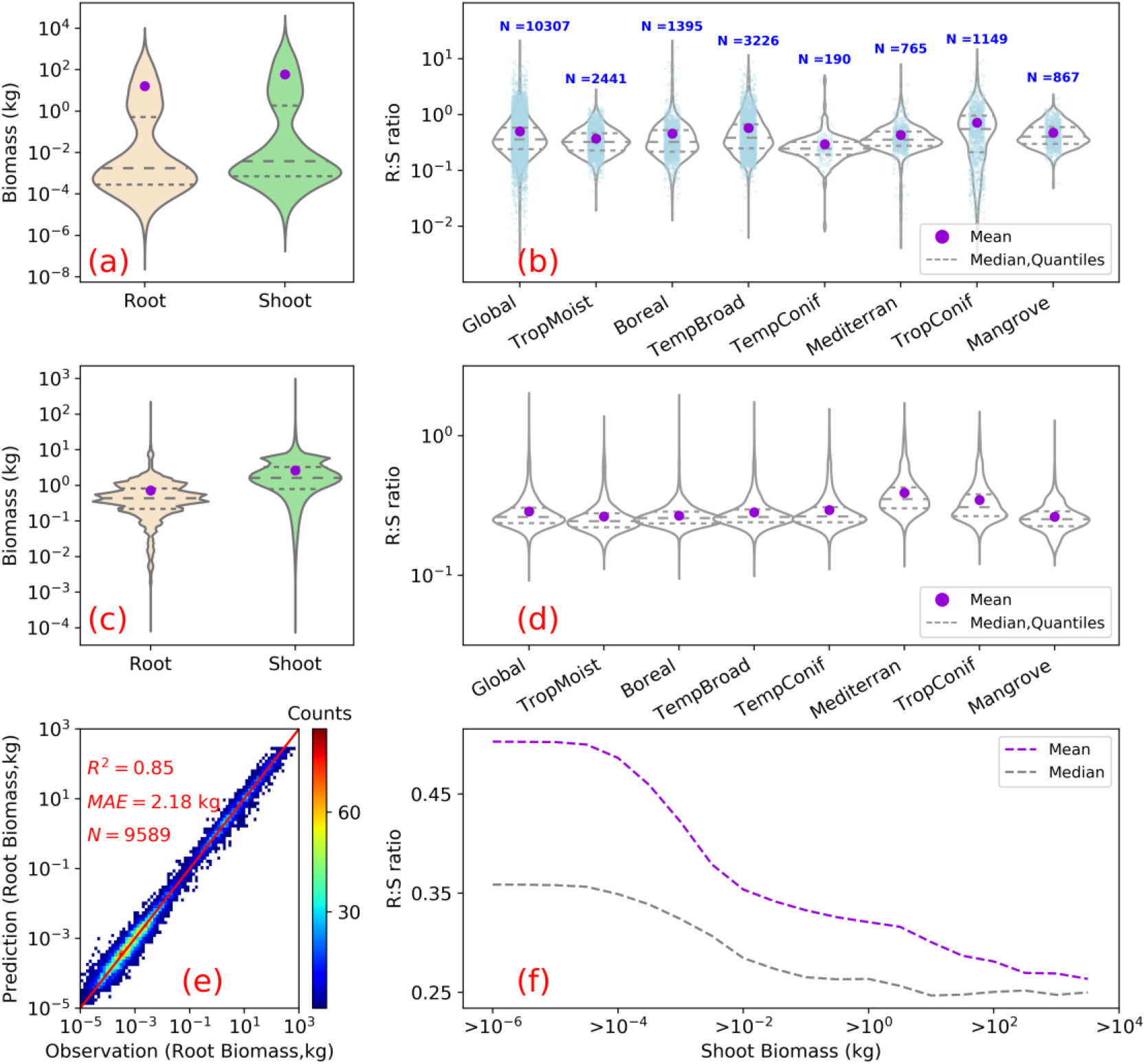
Root biomass and root shoot ratio (*R:S*). (a) and (b) show as violin plots the distribution of root and shoot biomass (in unit of kg/plant) and *R:S* ratios in the raw data used for upscaling. (c) and (d) are the distributions of model predicted root biomass from this study, of above-ground biomass used for the predicting, and of modelled *R:S* ratios at the global and biome scales. (e) is a heat plot of observed vs. predicted root biomass in kg of root per individual woody plant. (f) shows the mean (purple) and median (grey) *R:S* as a function of shoot biomass from observations. A shift of the shoot biomass towards a larger size ((a), (c)) results in a smaller predicted mean *R:S* at the global scale ((b),(d)) (see Supplementary Table 4 for exact values) as the mean *R:S* is size dependent (f). *R*^2^ is the coefficient of determination, *MAE* is the mean absolute error and *N* is the number of samples. TropMoist: tropical moist forest; Boreal: boreal forest/taiga; TempBroad: temperate broadleaf and mixed forest; TempConif: temperate coniferous forest; Mediterran: Mediterranean forests, woodlands and scrub; TropConif: tropical and subtropical coniferous forest; and Mangrove forest: mangrove forest. Note that the scales of y-axis are different between (a) and (c), (b) and (d). Model training and prediction were conducted on filtered data with *R:S* falling between the 1^st^ and 99^th^ percentiles and shoot biomass matching the range derived from GlobBiomass-AGB^6^ to reduce impacts from outliers.

We then analysed the dominant factors explaining spatial variations of root biomass and *R:S* (see Methods). Broadly speaking, locations with small trees, low precipitation, strong aridity, deep water table depth, high acidity, low bulk density, low base saturation and low cation exchange capacity are more likely to have higher fractional root biomass (Figure 3). In line with the allometric theory, shoot biomass emerged as the most important predictor of *R:S* and root biomass, as given by the Spearman correlation analysis shown in Figure 3, and partial importance plots (Supplementary Figures 7, 8, 9). Water related variables (precipitation, water table depth, aridity and vapor pressure) also emerged as important predictors in explaining *R:S* patterns (Figure 3)^20^, with trees and woody plants in dry regions generally having higher *R:S* (Supplementary Tables S3, S4), and with stronger dependence on precipitation when it is small and on water table depth when it is deep. Temperature is slightly negatively correlated with *R:S* at the global scale, in line with Reich et al. (2014). However, the relationship between temperature and below-ground biomass is not consistent among biomes (Figure 3) and biomass size groups (Supplementary Figures 7, 8, 9). The relationship between total soil nitrogen and root biomass is negative when soil nitrogen content is below 0.1-0.2 % (Supplementary Figure 7, 8, 9). Root biomass and *R:S* generally increases with soil alkalinity (Figure 3, Supplementary Figures 7, 8, 9). Low pH is toxic to biological activities and roots, especially fine roots are sensitive to soil acidification, as revealed by a recent meta-analysis^32^. Our results also indicate overall positive correlations between CEC, BS and R:S, but the processes that may account for these correlations are less clear from literature. Age has been shown to be important for R:S^33^. How age regulates *R:S* remains elusive, with studies showing positive^34^, slightly negative^35^ or no relationship^36^ between *R:S* and age. Including forest age (see Methods: Preparing predicting variables) as a predictor only marginally improved our model prediction (see SI for details). It is likely that shoot biomass partially accounts for age information and the quality of the global forest age data might also affect the power of this variable in improving root biomass predictions.

**Figure 3.**
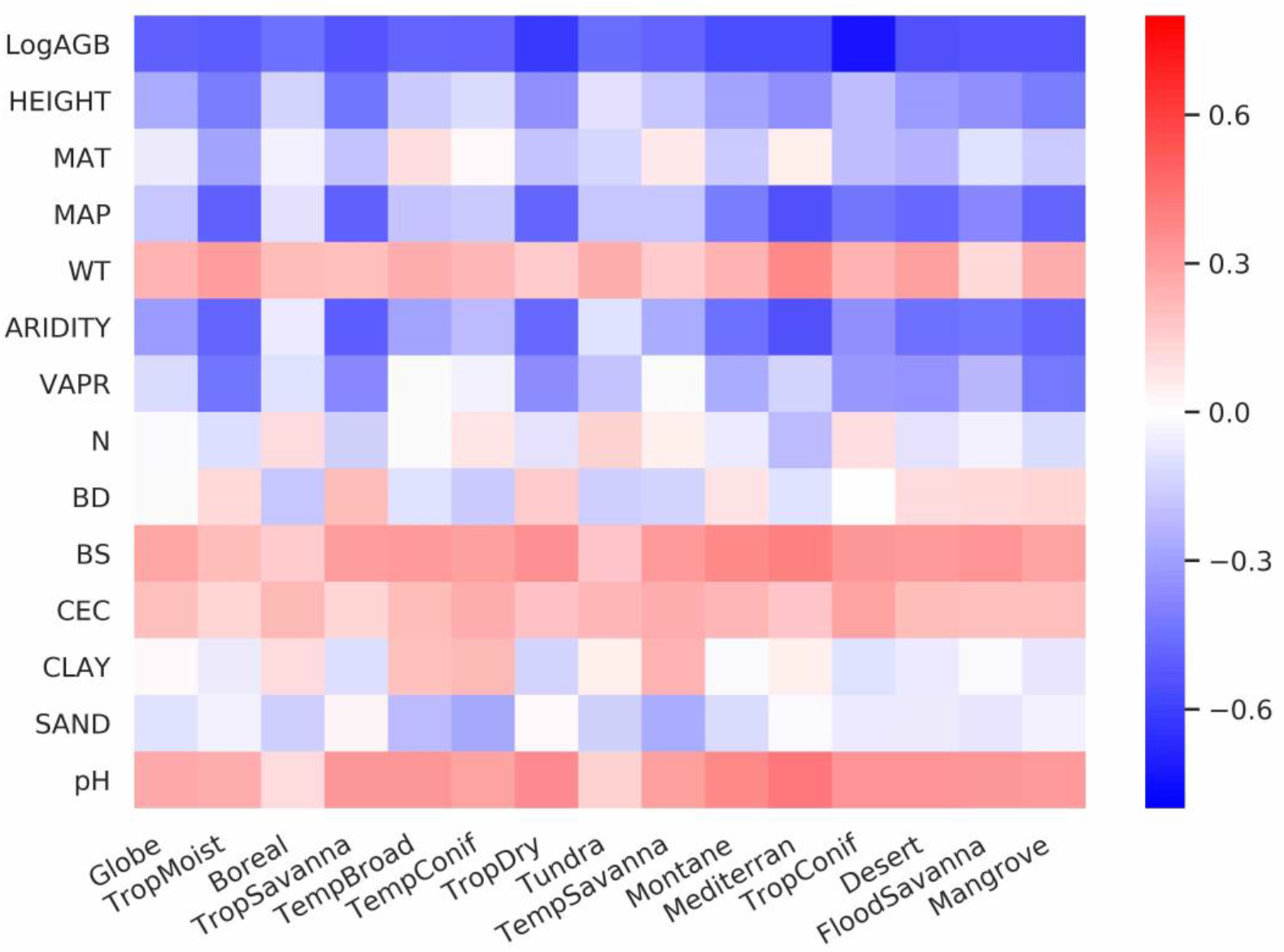
Spearman rank correlations between predicting variables and log-transferred *R:S*. Spearman coefficients are shown at both the global and biome scales for LogAGB: the logarithm of shoot biomass with base 10; HEIGHT: plant height; MAT: mean annual temperature; MAP: mean annual precipitation; WT: water table depth; ARIDITY: the aridity index; VAPR: water vapor pressure; N: soil nitrogen content; BD: soil bulk density; BS: soil base saturation; CEC, soil cation exchange capacity; CLAY: soil clay content; SAND: soil sand content; and pH: soil pH. From left to right, biomes are ordered descendingly according to their forest areas (Supplementary Figure 2).

## Discussion

Our lower estimation of root biomass compared to earlier studies is attributable to differences in forest area (Supplementary Table 5), above-ground biomass density (Supplementary Table 5), root biomass measurement and upscaling methodology. For example, the forest area in temperate zones used in Jackson et al. (1997) was about one third higher than in this study, which partly explains their higher root biomass for this biome (Supplementary Table 5). Our lower values of root biomass compared to Saugier et al. (2001), Mokany et al. (2006) and Robinson (2007) are caused mainly by our lower above-ground biomass density and *R:S* (Supplementary Table 5). Shoot biomass density (AGB) of tropical zones is 70% lower in our study than in Robinson (2007) who used sparse plot data collected more than a decade ago (Supplementary Table 5, case S2), and this lower AGB explains 27-46% of our lower root biomass (Supplementary Tables 5, 6). On the other hand, lower biome average *R:S* explains 41-48% of our underestimation compared to Robinson (2007). To elucidate this difference, we calculated weighted biome average *R:S* ratios through dividing total biome level shoot biomass by root biomass (i.e., weighted mean *R:S*). These weighted mean *R:S* ranging between 0.19 and 0.31 across biomes (Supplementary Table 3) are generally smaller than the *R:S* values reported in previous studies, which were based on average ratios obtained from sparser data (Supplementary Table 5), despite the arithmetic mean *R:S* (without weighting by biomass) from woody plants located in tropical, temperate and boreal zones (Supplementary Table 4) from our database being close to those from Robinson (2007).

The common practice of estimating root biomass through an average *R:S* without considering the high spatial variability of biomass and this ratio^4^ is a source of systematic error, leading to overestimating the global root biomass for two reasons. Firstly, upscaling ratios through arithmetic averages (possibly weighted by the number of trees or area, but not accounting for the fine grained distribution of biomass) systematically overestimates the true mean *R:S* (see SI Arithmetic mean *R:S* section) because *R:S* is a convex negative function of S given by *R*: *S* ∝ *S*^*β*−1^ with *β* taking typical values of about 0.9 ^35,37,38^. This explains why high-resolution S data used to diagnose weighted mean *R:S* ratios in our approach give generally smaller values than using arithmetic means at the biome level (see also Supplementary Tables 3 and 4). Secondly, available measurements tend to sample more small woody plants than big trees compared to real world distributions, because small plants are easier to excavate for measuring roots (see Figure 2a, 2c) but smaller plants tend to have larger *R:S* (Figure 2e). This sampling bias shifts the *R:S* towards larger values when using the mean from all samples in current databases. Our RF approach uses these data for training but in the upscaling, it accounts for realistic distributions of plant size. We further verified that our upscaled *R:S* ratios are robust to sub-sampling the training data in observed distributions, so that the bias of training data towards small plants does not translate into a bias of upscaled results.

The upscaling approach using allometric equations should also tend to overestimate the global root biomass due to the curvature of these allometric functions (see SI Allometric upscaling section). The global forest root biomass ranges between 154 – 210 Pg when root biomass was upscaled through different allometric equations collected from literature and fitted to our database (Supplementary Table 7), generally larger than from the RF mapping. Excluding the under-sampling issue in root biomass measurement, the global root biomass is likely to be smaller than when applying the allometric equation to the spatial average of shoot biomass (Supplementary Figures 10,11,12,13). Thus, future *in-situ* characterization of size structure across the world’s forests (see SI Allometric upscaling section) would greatly improve root biomass quantification.

An accurate spatially explicit global map of root biomass helps to improve our understanding of the Earth system dynamics by facilitating fundamental studies on resource allocation, carbon storage, plant water uptake, nutrient acquisition and other aspects of biogeochemical cycles. For example, the close correlation (correlation coefficient: 0.8) between root biomass and rooting depth^39^ at the global scale and the importance of root in plant water uptake and transpiration reflect close interactions between vegetation and hydrological cycles. The quest for drivers that affect allocation and consumption of photosynthetic production is a major focus of comparative plant ecology and evolution, as well as the basis of plant life history, ecological dynamics and global changes^11^. Turnover time and allocation are two key aspects that contribute to large uncertainties in current terrestrial biosphere model predictions^40,41^. Our root biomass map does not provide data on turnover or allocation, but an outcome on their aggregated effects. Future studies combining the root biomass map with upscaled root turnover data could shed light on the allocation puzzle. The growth of the fast turnover part of root, mostly fine root, and leaf are highly linked. If we assume an annual turnover of leaf and fine root, a preliminary estimation of average forest fine root biomass (from leaf biomass) reaches 6.7-7.7 Pg (see Supplementary Information: Preliminary estimation of fine root biomass). Despite being a small portion and highly uncertain, fine roots are temporally variable and functionally critical in ecosystem dynamics. Future studies on global distribution and temporal dynamics of fine roots are valuable. Considering specific biomes, tropical savannas would benefit from better root biomass estimation due to its large land area, and in tropical dry forests, field measurements of root and shoot biomass are needed to refine root biomass quantifications.

## Methods

### Overview

Our global mapping of root biomass relies on a predicting model based on a machine learning algorithm that is fitted to a large number of ground field measurements. Root biomass was upscaled as a function of shoot biomass, tree height, age, species, land management, topography, edaphic and climate variables. The process takes three major steps (Supplementary Figure 1). The first step is to collect field measurements, and observations of auxiliary variables such as tree height, age, species and management status (see sections field measurements and preparing predicting variables below). In a second step, we compared the allometric upscaling and tested three machine learning techniques, the random forest (RF), the artificial neural networks (ANN) and multiple adaptive regression splines (MARS) through 47 input variables. The best predicting model with the minimum number of predictors and with the lowest mean absolute error (MAE) and highest R-squared value (R^2^) was selected through cross-validation (see section Building predicting models below). The next step was to generate a 1 km global root biomass map by running the best predicting model on spatially-explicit gridded fields of model inputs. The model outputs were initially expressed as root biomass in unit of weight per individual woody plant and were then mapped into root biomass per unit area using tree densities (the number of trees per unit area)^25^. The uncertainty of the mapping and the importance of the model inputs were analysed in detail as explained below.

### Field measurements

Our dataset was compiled from literature and existing forest biomass structure or allometry databases^42 20,33,43^. We included studies and databases that reported georeferenced location, root biomass and shoot biomass. For example, Ref^44^ is not included due to lack of georeferenced location and Ref^45^ in not used as we also need measurements of other plant compartments like shoot biomass. Repeated entries from existing databases were removed. One of the databases^42^ reported data on woody plants which also include shrub species. We kept the shrub data partly because the remote sensing products we used to generate our root map do not clearly separate trees from shrubs. Around 82% of the extracted entries also recorded plant height and management status. Height was identified as an important predictor in our model assessment, and entries were discarded when height was missing (18% of data). As woody plant age was reported in 19% of the entries only, the values of this variable was determined from another source of information, i.e. from a composite global map introduced in the next section. Species names were systematically reported, but biotic, climatic, topographic and soil information were missing for a substantial proportion of entries and values of these variables were thus extracted from independent observation-driven global maps as explained in the next section. Our final dataset includes biomass measurements collected in 494 different locations from 10307 individual plants, which cover 465 species across 10 biomes as defined by The Nature Conservancy^22^ (Supplementary Figure 2; Supplementary Data).

### Preparing predicting variables

We used 47 predictors that broadly cover 5 categories: vegetative, edaphic, climatic, topographic and anthropogenic (Supplementary Table 1). Vegetative variables include shoot biomass, height, age, maximum rooting depth, biome class and species. Edaphic predictors cover soil bulk density, organic carbon, pH, sand content, clay content, total nitrogen, total phosphorus, Bray phosphorus, total potassium, exchangeable aluminium, cation exchange capacity, base saturation (BS), soil moisture and water table depth (WT). Climatic predictors are mean annual temperature (MAT), mean annual precipitation (MAP), the aridity index that represents the ratio between precipitation the reference evapotranspiration, solar radiation, potential evapotranspiration (PET), vapor pressure, cumulative water deficit (CWD=PET - MAP), wind speed, and mean diurnal range of temperature (BIO2), isothermality (BIO2/BIO7) (BIO3), temperature seasonality (BIO4), max temperature of warmest month (BIO5), min temperature of coldest month (BIO6), temperature annual range (BIO7), mean temperature of wettest quarter (BIO8), mean temperature of driest quarter (BIO9), mean temperature of warmest quarter (BIO10), mean temperature of coldest quarter (BIO11), precipitation of wettest month (BIO13), precipitation of driest month (BIO14), precipitation seasonality (BIO15), precipitation of wettest quarter (BIO16), precipitation of driest quarter (BIO17), precipitation of warmest quarter (BIO18), precipitation of coldest quarter (BIO19). The topographic variable is elevation and we take the management status (managed or not) as the anthropogenic predictor. All references are given in Supplementary Table 1.

To derive the shoot or above-ground biomass (AGB) per tree (in unit of weight per tree), we combined the GlobBiomass-AGB satellite data product^24^ (in unit of weight per unit area) with a tree density map (number of trees per unit area)^25^. The GlobBiomass dataset was based on multiple remote sensing products (radar, optical, LiDAR) and a large pool of *in-situ* observations of forest variables^6,46^. The original GlobBiomass-AGB map was generated at 100 m spatial resolution; for this study, the map was averaged into a 1 km pixel by considering only those pixels that were labeled as forest ^6^. A pixel was labeled as forest when the canopy density was larger than 15% according to Hansen et al. (2013)’s dataset (Hansen2013) averaged at 100 m. The 1-km resolution global tree density map was constructed through upscaling 429,775 ground-based tree density measurements with a predictive regression model for forests in each biome^25^. The forest canopy height map took advantage of the Geoscience Laser Altimeter System (GLAS) aboard ICESat (Ice, Cloud, and land Elevation Satellite). Forest definitions are slightly different among these three maps. Forest area of the tree density map was based on a global consensus land cover dataset that merges four land cover products ^47^, which gave an equal total tree count as the Hansen et al. (2013) land cover ^25^. The canopy height map used the Globcover land cover map^48^ as reference to define forest land. We took Hansen2013 with a 15% canopy cover threshold as our base forest cover map. We approximated the missing values in tree density and height (due to mismatches in forest cover) by the mean of a 5×5 window that is centered on the corresponding pixel. We quantified the potential impact of mismatches in forest definition by looking into two different thresholds: 0% and 30%.

We merged several regional age maps to generate a global forest age map. The base age map was derived from biomass through age-biomass curve similarly as conducted in tropical regions in ref.^49^ This age map does not cover the northern region beyond 35 N. We filled the missing northern region with a North American age map ^50^ and a second age map covers China^51^. Remaining missing pixels were further filled with the age map derived from MODIS disturbance observations. For the final step, we filled the remaining pixels with the GFAD V1.1 age map^49^. GFAD V1.1 has 15 age classes and 4 plant functional types (PFTs). We choose the middle value of each age class and estimated the age as the average among different PFTs.

Detailed information of all ancillary variables is listed in Supplementary Table 1. To stay coherent, we re-gridded each map to a common 1 km × 1 km grid through the nearest neighbourhood method.

### Building predicting models

We investigated the performance of the allometric scaling and three non-parametric models: RF, ANN and MARS. Allometric upscaling relates root biomass to shoot biomass in the form of *R* ∝ *S*^*β*^. RF is an ensemble machine learning method that builds a number of decision trees through training samples^52^. A decision tree is a flow-chart-like structure, where each internal (non-leaf) node denotes a binary test on a predicting variable, each branch represents the outcome of a test, and each leaf (or terminal) node holds a predicted target variable. With a combination of learning trees (models), RF generally increases the overall predicting performance and reduces over-fitting. ANN computes through an interconnected group of nodes, inspired by a simplification of neurons in a brain. MARS is a non-parametric regression method that builds multiple linear regression models across a range of predictors.

Tree shoot biomass from the *in-situ* observation data spans a wider range than shoot biomass per plant derived from global maps (1×10^−7^ to 8800 vs. 7.9×10^−5^ to 933 kg/plant). To reduce potential mapping errors, we selected training samples with shoot biomass between 5×10^−5^ and 1000 kg/plant. The medians and means of shoot biomass, root biomass and *R:S* from the selected training samples are similar as that from the entire database. Also, to reduce the potential impact of outliers, we analyzed samples with *R:S* falling between the 1^st^ and 99^th^ percentiles, which consists of 9589 samples with *R:S* ranging from 0.05 to 2.47 and a mean of 0.47 and a median of 0.36. Sample filtering slightly deteriorated model performance and had minor impact on the final global root biomass prediction (145 from whole samples vs.142 Pg from filtered data). We chose root biomass as our target variable instead of *R:S* because big and small trees contribute equally to *R:S* while big trees are relatively more important in biomass quantification. In our observation database, we have more samples being small woody plants. A predicting model with an overall good performance will not guarantee a good prediction on woody plants with higher biomass. We, furthermore split the *in-situ* measured shoot biomass into three groups, namely with shoot biomass smaller than 0.1, between 0.1 and 10, and larger than 10 kg/plant. The rationale behind this splitting is: (1), the distribution of *in-situ* measured woody shoot biomass (Figure 2); (2), empirical evidence showing the shift of root shoot allometry with tree size^44 20^; (3), a better performance on independent validation samples through numerous combinations of splitting trials; (4), tests through weighting samples or resampling samples (e.g., over-sampling using Synthetic Minority Over-sampling Technique) gave no better performance.

Model performances were assessed by 4-fold cross-validation using two criteria: the mean absolute error (MAE), the R-squared value (R^2^). MAE quantifies the overall error while R^2^ estimates the proportion of variance in root biomass that is captured by the predicting model. We favored the model with a smallest MAE, a highest R^2^ and with minimum number of predictors. For non-parametric models, starting from a model with all 47 predictors, we sequentially excluded predictors that did not improve model performance one after another. The order of predictor removing is random. After a combination of trials, the best model is from RF and the final set of predictors include shoot biomass, height, soil nitrogen, pH, bulk density, clay content, sand content, base saturation, cation exchange capacity, vapor pressure, mean annual precipitation, mean annual temperature, aridity and water table depth.

### Generation of the global root biomass map

We assumed shoot size and other selected predictors to be drivers of root biomass. Building upon a large set of samples with each field measurement being an outcome of complex local interactions (including within-vegetation competition), we implicitly accounted for sub-pixel variability (e.g., resource competition and responses to environmental conditions) on allometry. Biome class and species were excluded from the pool of predicting variables because they did not improve model performance. We combined the RF model with global maps of selected predicting variables to produce the root biomass map which has a unit of weight per tree. This map was multiplied by tree density at 1-km resolution to obtain the final root biomass map with a unit of weight per area (Supplementary Figure 1).

### Uncertainty quantification

We estimated the overall uncertainty of the root biomass estimates through quantifying relative errors caused by predictors at the 1-km resolution, predicting errors associated with RF given correct predicting variables, and errors from upscaling root biomass per tree to root biomass per unit area.

#### Predictor errors (η_pred_)

We collected 8 additional global predictor datasets (3 shoot biomass, 2 soil and 3 climate datasets) (Supplementary Table 2). We carried out 8 sets of additional predictions replacing the predictors by each of these additional data maps and calculated the standard deviation among 8 predictions for each pixel. The overall predictor errors were expressed in a relative term, that is, the ratio between the standard deviation and the standard prediction (with the GlobBiomass-AGB and other predictors listed in Supplementary Table 1) for each pixel.

#### RF errors (η_RF_)

The performance of machine learning models is frequently verified through the independent test samples. We carried out 4-fold cross-validation. The RF error is quantified as the relative error (the standard deviation divided by the mean) from 4-fold predictions.

#### Upscaling errors (η_up_)

Upscaling the root biomass from per tree to per area relies on the tree density map. The upscaling error is set as the relative uncertainty of tree density^25^.

At last we propagated these relative errors across the entire root biomass quantification processes assuming these three errors were random and independent. So the errors were assumed to be uncorrelated and the covariation were assumed to be 0. The overall relative errors at the pixel level was calculated through,

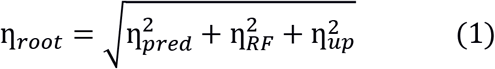

Uncertainty at the global or biome scale (*σ*_*biome*_) is quantified through expanding calculating area and propagating the relative errors at the pixel level,

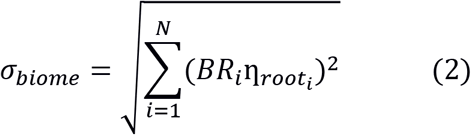

where BR is the total root biomass (in unit of weight) in each forested pixel and N is the number of pixels within biome boundaries (or all forested pixels when calculate the global total). η_*root*__*i*_ is the relative uncertainty in quantifying root biomass for the *ith* pixel.

### Relative importance of predicting variables

The impact of predictors on predicting *R:S* was estimated through the Spearman’s rank-order correlation at both the global and biome scales. We log-transformed the *R:S* and shoot biomass before standardizing these datasets. Partial dependence plot^53^ tells the marginal effect of one predictor have on root biomass from a machine learning model, and serves as a supplement to the Spearman correlation.

## Acknowledgements

Y.H., D.S.G and P.C. received support from the European Research Council Synergy project SyG-2013-610028 IMBALANCE-P and P.C. and Y.H. from the ANR CLAND Convergence Institute. H.Y., D.S. and M. S. were funded through the ESA Climate Change Initiative BIOMASS project. Collecting Russian data were supported by The Russian Science Foundation (project no. 19-77-30015).

## Author contributions

Y.H. and P.C. designed this study. Y. H., P.C., M.S., J.C and D.S. collected the data. D.M., P.C., M.S., J.C., Y.C. and Y.H. discussed analyzing methods. Y.H. conducted the analysis and drafted the manuscript. All authors discussed the results and contributed to the manuscript.

## Code availability

Calculations were conducted through Python 2.7.15 and ferret 6.72. The code is available upon request.

## Data availability

The datasets generated in this study are available from the corresponding author on reasonable request.

