## Supplemental material for "A global map of root biomass across the world’s forests"

**Supplementary Figures and Tables**

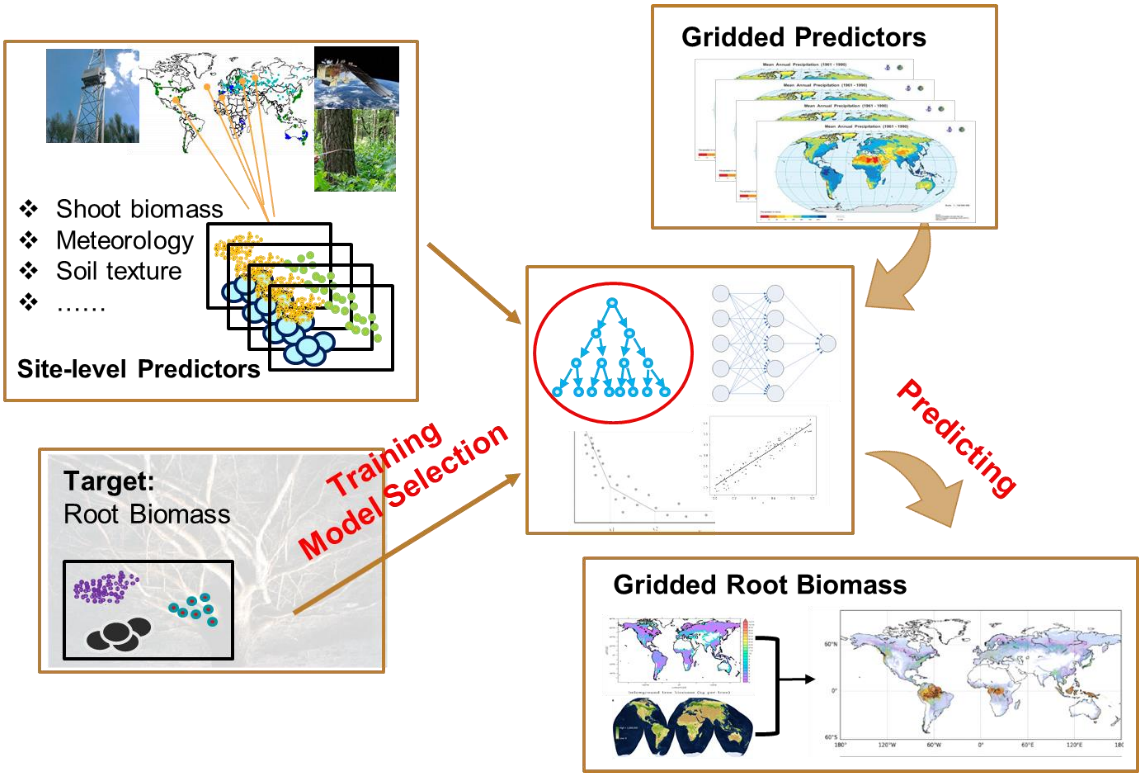


Figure 1. Procedures of root biomass mapping at the 1-km resolution. Root biomass mapping takes 3 major steps. Step 1: compile field measurements and prepare global gridded predictors; Step 2: train the model with data from Step 1 and select the model with best performance; and Step 3, map root biomass with selected model from Step 2 and gridded predictors from Step 1. We split the data into 3 size categories and selected among 47 predictors through 4 modeling methods (the allometric equation, the random forest, the artificial neural networks and multiple adaptive regression splines). The final root biomass map with a unit of weight per area is created through combining the predicting results (in unit of weight per individual tree) with the tree density (number of trees per area).

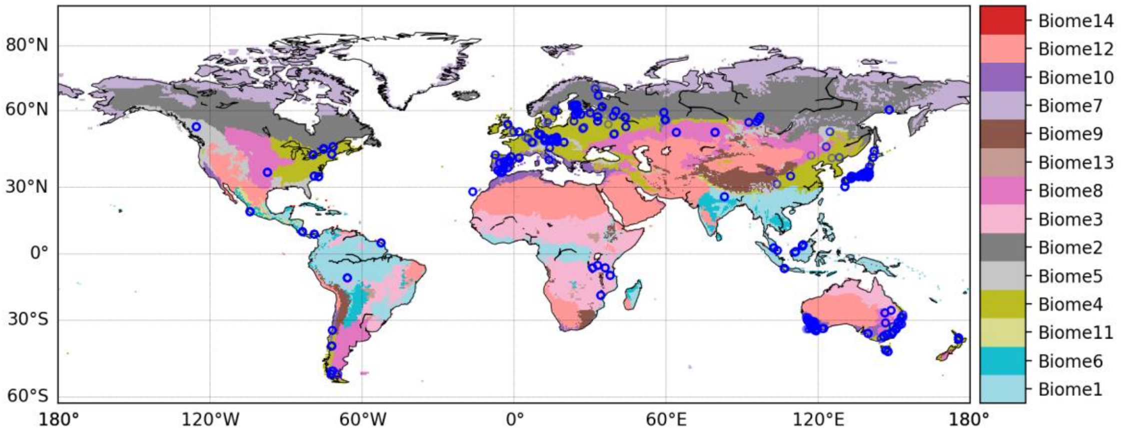


Figure 2. Geographical distribution of observation sites (blue circles) and biome classes from The Nature Conservancy[1]. Numbers after Biome from the legend are ordered incrementally with decreasing forest area of each biome (Table 3). Biome 1: tropical moist forests; Biome 2:boreal and taiga forests; Biome 3: tropical and subtropical grasslands, savannas and shrublands; Biome 4: temperate broadleaf and mixed forests; Biome 5: temperate coniferous forests; Biome 6: tropical dry forests; Biome 7: tundra; Biome 8: temperate grasslands, savannas and shrublands; Biome 9: montane grasslands and shrublands; Biome 10: Mediterranean forests, woodlands and scrubs; Biome 11: tropical and subtropical coniferous forests; Biome 12: deserts and xeric shrubland; Biome 13: flooded grasslands, savannas; and Biome 14: mangroves.

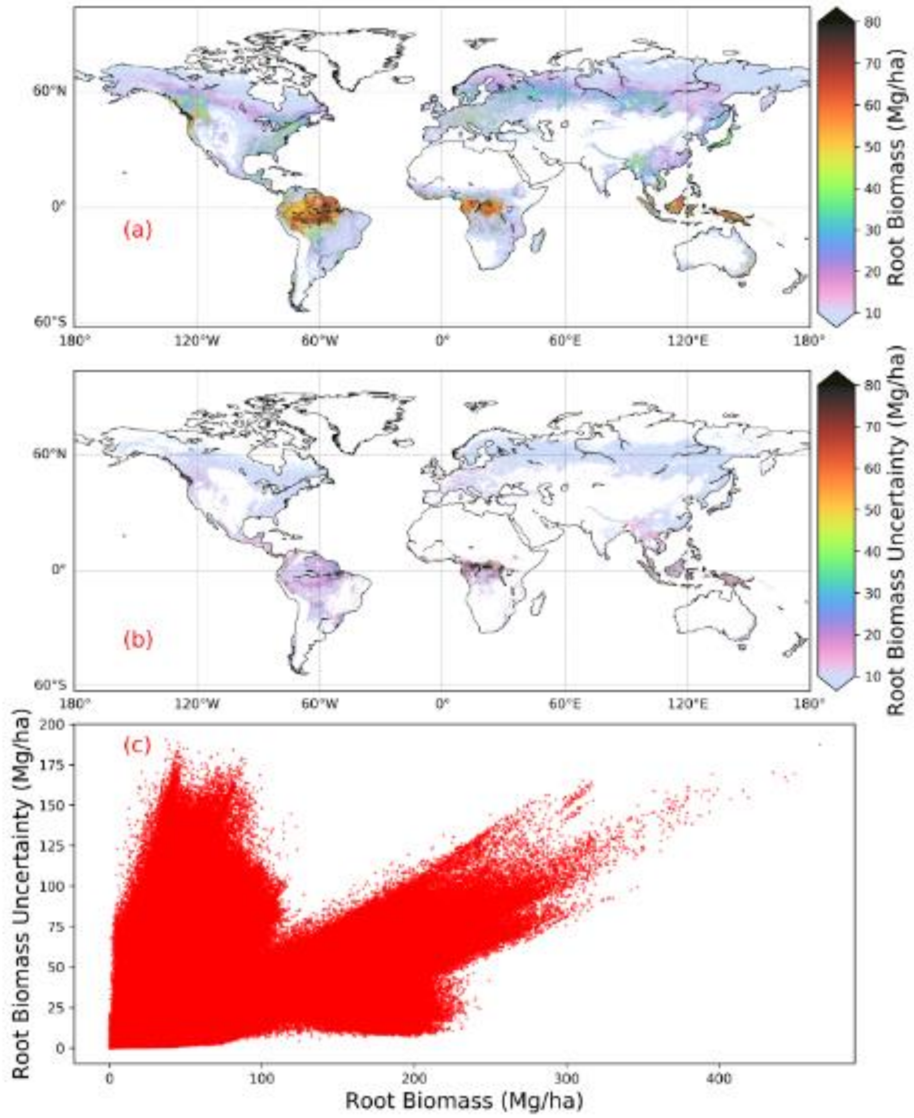


Figure 3. Spatial distribution of root biomass (a) and mapping uncertainty (b) at 1 km spatial resolution, and the scatter plot of root biomass vs. mapping uncertainty (c).

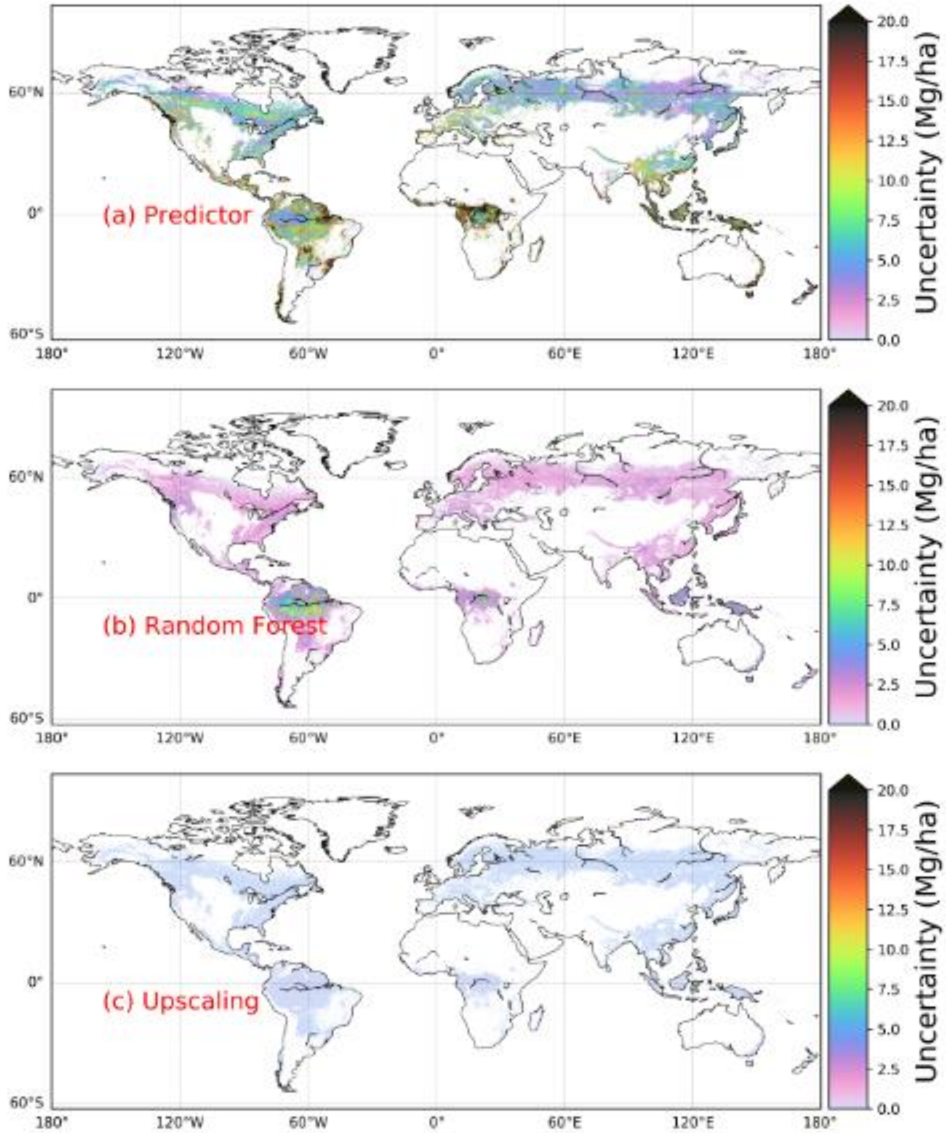


Figure 4. Contributions of predictor uncertainty (a), random forest model (b) and upscaling (c) to the overall uncertainty in root biomass estimation.

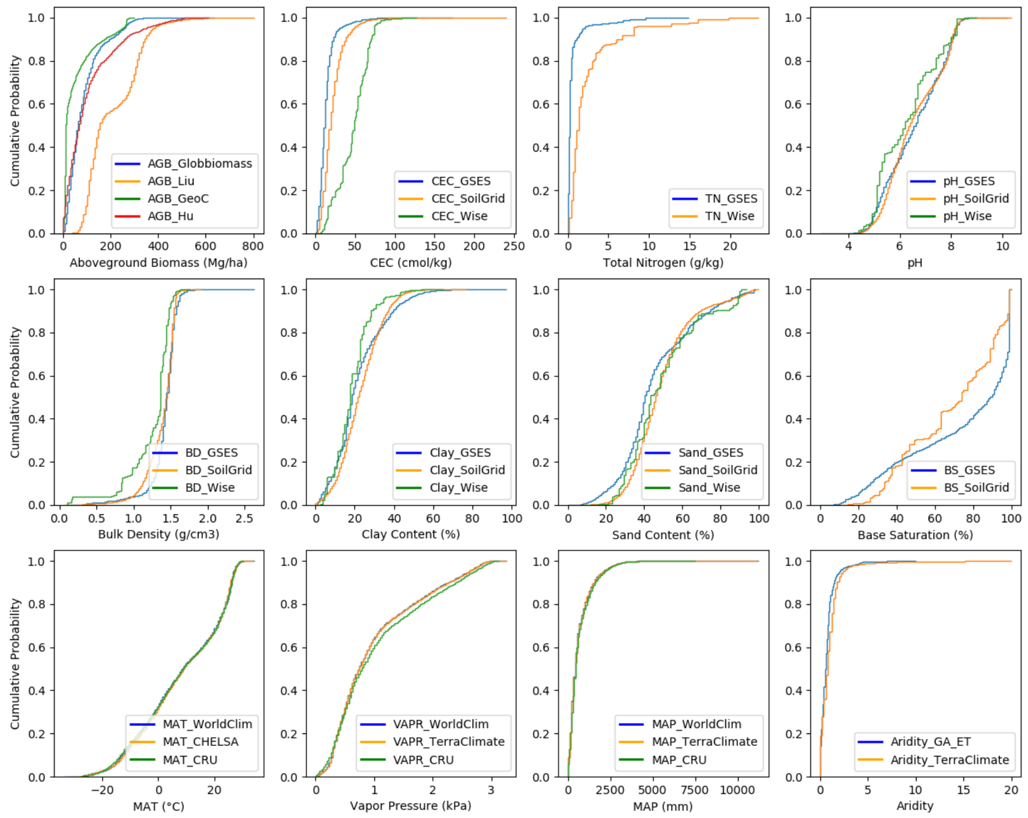


Figure 5. Cumulative distributions of predictors. Each panel corresponds to one predictor used in quantifying the contribution of predictors to uncertainty in root biomass mapping (Figure 4a). Different colors indicate different sources for each predictor. Detailed information of data sources is provided in Tables 1, 2.

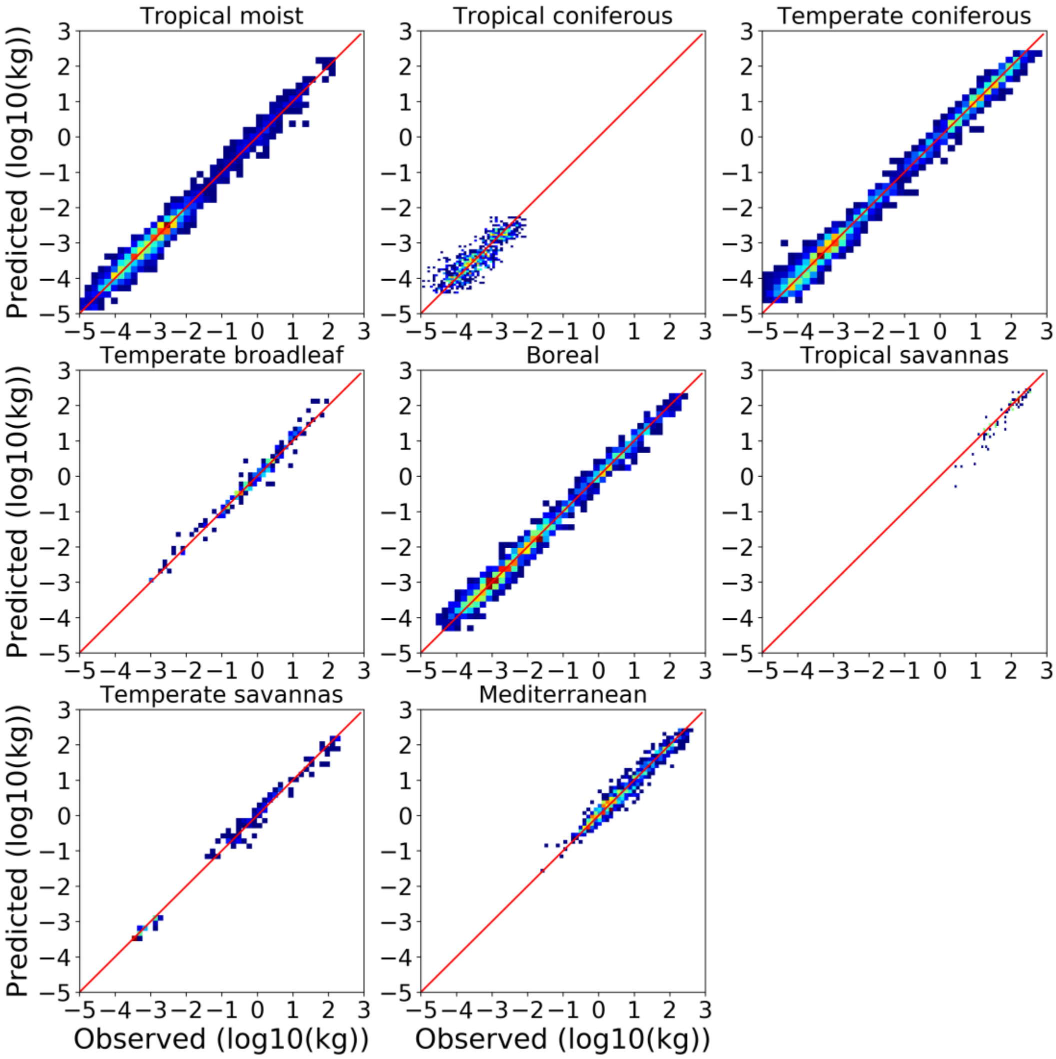


Figure 6. Heat plots of predicted root biomass vs. observation at the biome level. Biome classification is from The Nature Conservancy[1] and is shown in Figure 2.

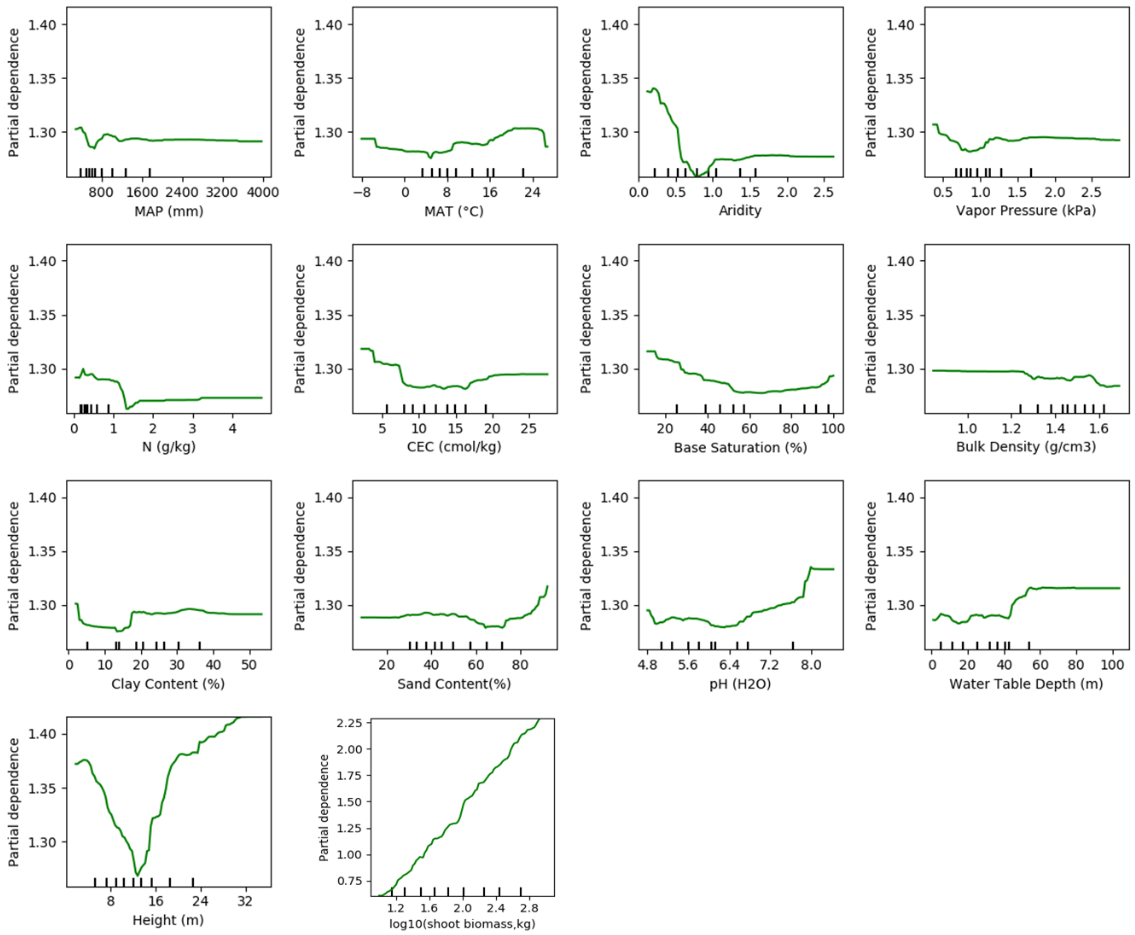


Figure 6. Partial dependence plot showing the dependence of root biomass on predictors for woody plant with shoot biomass > 10 kg. 10 kg is one threshold based on which we split our datasets for a best model performance (see Methods). Note the y-axis of the last panel (shoot biomass) is different from other predictors.

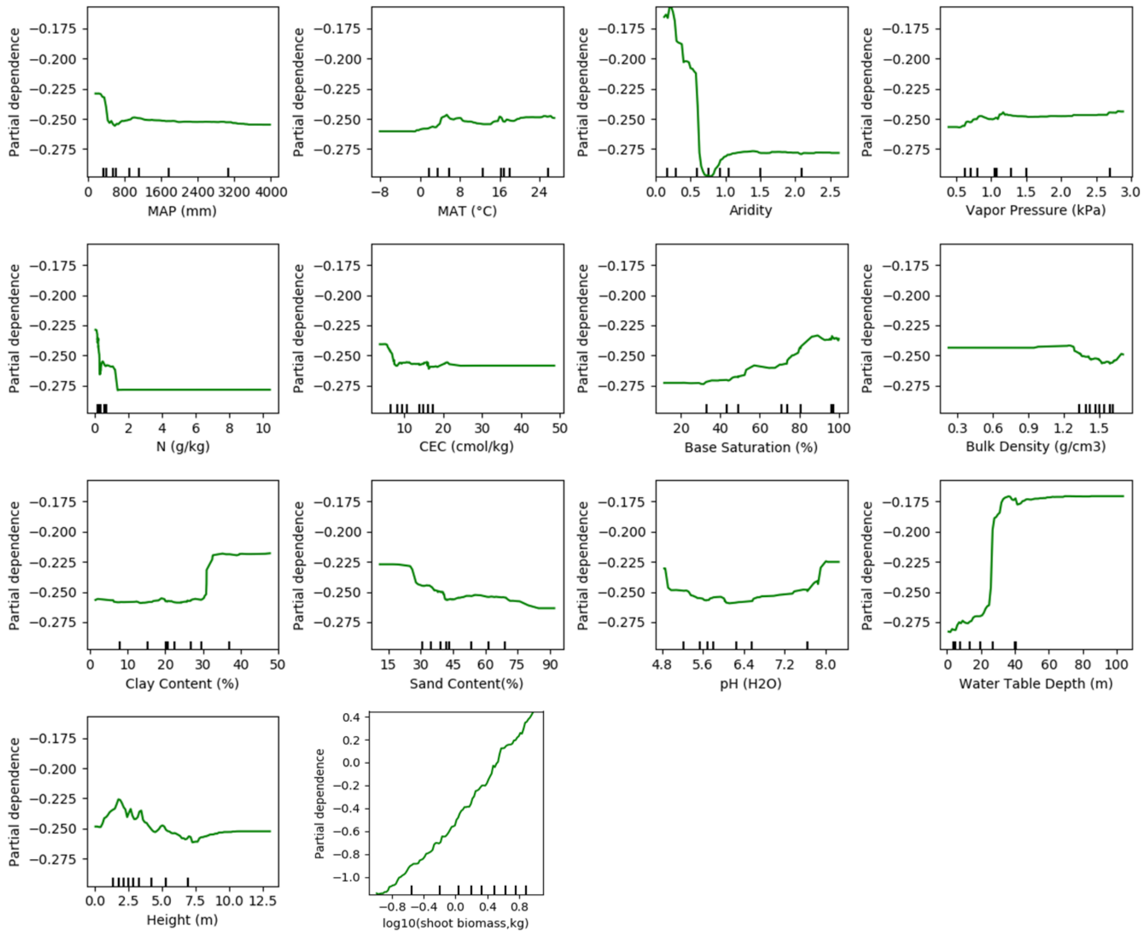


Figure 7. Partial dependence plot showing the dependence of root biomass on predictors for woody plant with shoot biomass between [0.1 10] kg. 0.1 and 10 kg are thresholds based on which we split our datasets for a best model performance (see Methods). Note the y-axis of the last panel (shoot biomass) is different from other predictors.

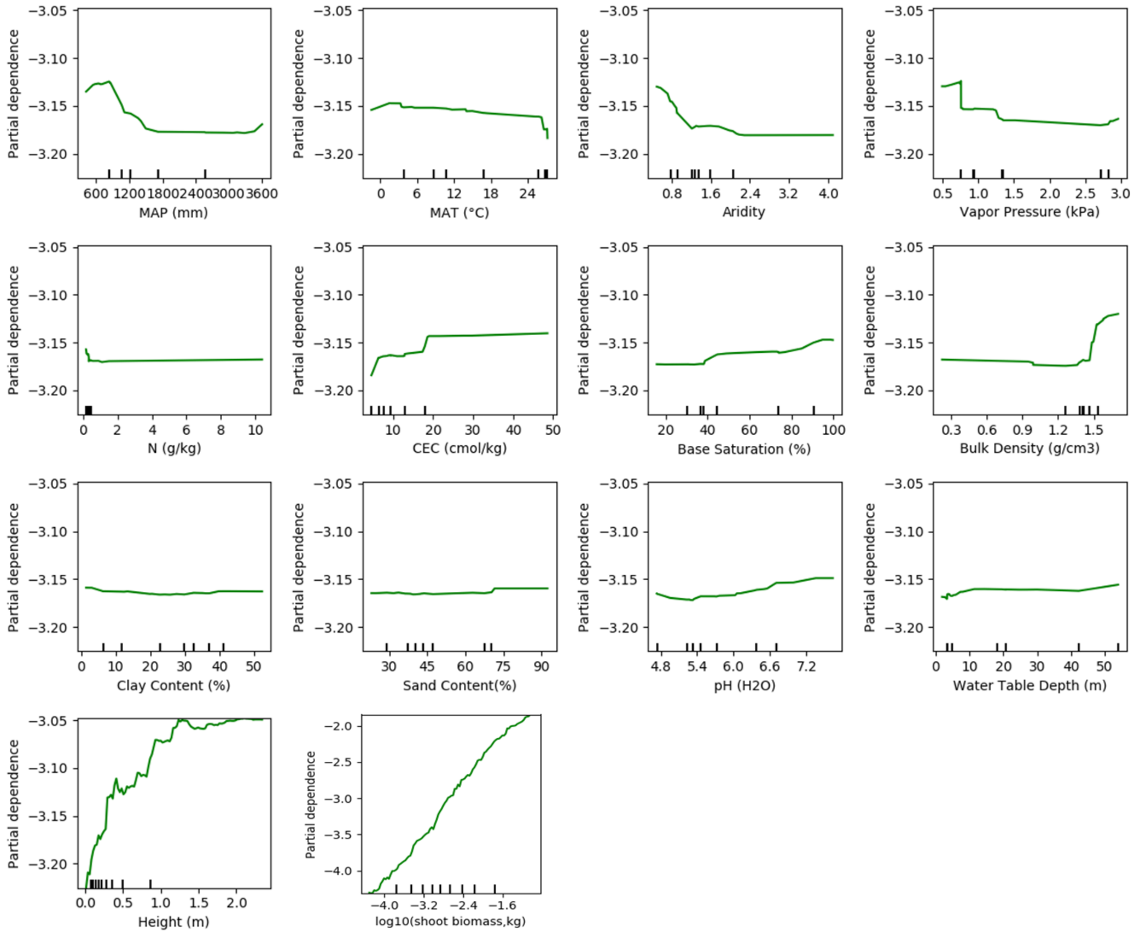


Figure 8. Partial dependence plot showing the dependence of root biomass on predictors for woody plant with shoot biomass smaller than 0.1 kg. 0.1 kg is one threshold based on which we split our datasets for a best model performance (see Methods). Note the y-axis of the last panel (shoot biomass) is different from other predictors.

Table 1. The source, unit, category, resolution, time coverage and reference of gridded global datasets used in building training model and predicting root biomass. BIO2-11 and BIO13-19 corresponds to Bioclimatic variables from WorldClim version 2. All datasets were accessed in February 2019.

| **Name** | **Source** | **Unit** | **Type** | **Res** | **Time** | **Reference** |
|---|---|---|---|---|---|---|
| Age | Mixed | year | Biological | 1km | Current | See Methods for details |
| Maximum Rooting Depth | GSES | m | Biological | 1km | Current | http://globalchange.bnu.edu.cn/research/soilw |
| Biome | The nature conservancy | | Biological | 1km | Current | http://maps.tnc.org/gis_data.html |
| Height | Simard | m | Biological | 1km | Current | https://webmap.ornl.gov/wcsdown/dataset.jsp?ds_id=10023 |
| Aboveground biomass density | GlobBiomass | Mg/ha | Biological | 1km | Current | http://globbiomass.org/wp-content/uploads/GB_Maps/Globbiomass_global_dataset.html |
| Tree density | Crowther | per ha | Biological | 1km | Current | https://elischolar.library.yale.edu/yale_fes_data/1/ |
| Rooting depth | Fan | m | Biological | | Current | https://wci.earth2observe.eu/thredds/catalog/usc/root-depth/catalog.html |
| Bulk Density | GSES | g/cm$^3$ | Soil | 1km | Current | http://globalchange.bnu.edu.cn/research/soilw |
| Soil Organic Matter | GSES | % of weight | Edaphic | 1km | Current | http://globalchange.bnu.edu.cn/research/soilw |
| Soil pH | GSES | | Edaphic | 1km | Current | http://globalchange.bnu.edu.cn/research/soilw |
| Soil Sand | GSES | % of weight | Edaphic | 1km | Current | http://globalchange.bnu.edu.cn/research/soilw |
| Soil Clay | GSES | % of weight | Edaphic | 1km | Current | http://globalchange.bnu.edu.cn/research/soilw |
| Total Nitrogen | GSES | % of weight | Edaphic | 1km | Current | http://globalchange.bnu.edu.cn/research/soilw |
| Total Phosphorus | GSES | % of weight | Edaphic | 1km | Current | http://globalchange.bnu.edu.cn/research/soilw |
| Bray Phosphorus | GSES | ppm | Edaphic | 1km | Current | http://globalchange.bnu.edu.cn/research/soilw |
| Total Potassium | GSES | % of weight | Edaphic | 1km | Current | http://globalchange.bnu.edu.cn/research/soilw |
| Exchangeable Aluminum | GSES | cmol/kg | Edaphic | 1km | Current | http://globalchange.bnu.edu.cn/research/soilw |
| Cation Exchange Capacity | GSES | cmol/kg | Edaphic | 1km | Current | http://globalchange.bnu.edu.cn/research/soilw |
| Base Saturation | GSES | % | Edaphic | 1km | Current | http://globalchange.bnu.edu.cn/research/soilw |
| Soil Moisture | ESA CCI | m3/m3 | Edaphic | 0.25° | Average 1982-2005 | https://www.esa-soilmoisture-cci.org/ |
| Water Table Depth | Fan2013 | m | Edaphic | 1km | Current | https://glowasis.deltares.nl/thredds/catalog/opendap/opendap/Equilibrium_Water_Table/catalog.html |
| Mean Annual Precipitation | WorldClim V2.0 | mm | Climatic | 1km | Average 1970-2000 | http://www.worldclim.org |
| Mean Annual Temperature | WorldClim V2.0 | °C | Climatic | 1km | Average 1970-2000 | http://www.worldclim.org |
| Aridity | GA-ET | | Climatic | 1km | Average 1970-2000 | https://figshare.com/articles/Global_Aridity_Index_and_Potential_Evapotranspiration_ET0_Climate_Database_v2/7504448/3 |
| Potential Evapotranspiration | GA-ET | mm | Climatic | 1km | Average 1970-2000 | https://figshare.com/articles/Global_Aridity_Index_and_Potential_Evapotranspiration_ET0_Climate_Database_v2/7504448/3 |
| Solar Radiation | WorldClim V2.0 | kJ/m2 /day | Climatic | 1km | Average 1970-2000 | http://www.worldclim.org |
| Vapor | WorldClim | kPa | Climatic | 1km | Average | http://www.worldclim.org |

| | | | | | | |
|---|---|---|---|---|---|---|
| Pressure | V2.0 | | | | 1970-2000 | |
| Cumulative Water Deficit | WorldClim V2.0 | mm | Climatic | 1km | Average 1970-2000 | PET - MAP |
| Wind Speed | WorldClim V2.0 | m/s | Climatic | 1km | Average 1970-2000 | http://www.worldclim.org |
| BIO2-11 | WorldClim V2.0 | | Climatic | 1km | Average 1970-2000 | http://www.worldclim.org |
| BIO13-19 | WorldClim V2.0 | | Climatic | 1km | Average 1970-2000 | http://www.worldclim.org |
| Elevation | SRTM30_PLUS v8 | m | Topographical | 1km | Average 1970-2000 | https://eatlas.org.au/data/uuid/80301676-97fb-4bdf-b06c-e961e5c0cb0b |

Table 2. Alterative global datasets for quantifying root biomass predicting uncertainty. All datasets were accessed in June 2019.

| Name | Variables | Res | Time | Reference |
|---|---|---|---|---|
| AGB_Hu | Shoot biomass | 1km | Current | Hu, et al. (2016) |
| AGB_Liu | Shoot biomass | 0.25° | 1993-2012 | Liu, et al. (2015) |
| AGB_GeoC | Shoot biomass | 0.01 | Current | GEOCARBON, https://www.bgc-jena.mpg.de/geodb/projects/Home.php |
| SoilGrid | CEC, Bulk density, Clay content, Sand content ,CEC, | 1km | Current | Hengl, et al. (2017) |
| WISE30 | Total nitrogen, pH, Bulk density, clay, sand, Base saturation, CEC, | 1km | Current | Batjes (2015) |
| CHELSA | MAT | 1km | Same as WorldClim | http://chelsa-climate.org/ |
| TerraClimate | Aridity, MAP, Vapor pressure | 4 km | Same as WorldClim | http://www.climatologylab.org/terraclimate.html |
| CRU_TS4.03 | Vapor pressure, MAP, MAT, aridity | 0.5° | Same as WorldClim | https://crudata.uea.ac.uk/cru/data/hrg/ |

Table 3. Land area, land area occupied by woody plants (forest area), shoot biomass, root biomass and weighted *R:S* ratio (total shoot biomass/total root biomass) at the biome and global scales. The biome classification is from The Nature Conservancy[1]. Forest area covers land with canopy cover > 15%[6]. Numbers after ± are uncertainties quantified in GlobBiomass[7] for shoot biomass and in this study for root biomass and *R:S* (see Methods).

| Biome number | Name | Land area ($10^6$ km$^2$) | Forest area ($10^6$ km$^2$) | Shoot biomass (Pg) | Root biomass (Pg) | Weighted *R:S* Ratio |
|---|---|---|---|---|---|---|
| 1 | Tropical moist | 19.8 | 15.6 | 295±103.5 | 71.7±22.3 | 0.24±0.11 |
| 2 | Boreal | 16 | 11.2 | 77.5±22.0 | 19.5±0.7 | 0.25±0.08 |
| 3 | Tropical savanna | 19.5 | 6.7 | 52±15.2 | 13.7±0.2 | 0.26±0.08 |
| 4 | Temperate broadleaf | 12.9 | 5.8 | 66±16.2 | 16.6±2.7 | 0.25±0.08 |
| 5 | Temperate coniferous | 4.4 | 2.5 | 32.2±7.8 | 8.2±1.5 | 0.25±0.08 |
| 6 | Tropical dry | 3.8 | 1.4 | 13.7±3.7 | 3.8±2.9 | 0.28±0.10 |
| 7 | Tundra | 8.0 | 0.9 | 3.9±0.7 | 1.1±1 | 0.28±0.06 |
| 8 | Temperate savanna | 9.6 | 0.7 | 4.7±0.8 | 1.4±0.1 | 0.30±0.06 |
| 9 | Montane | 5.2 | 0.5 | 4.3±1.0 | 1.3±0.2 | 0.30±0.09 |
| 10 | Mediterranean | 3.3 | 0.5 | 4.8±0.8 | 1.5±0. | 0.31±0.07 |

| | | | | | | |
|---|---|---|---|---|---|---|
| 11 | Tropical coniferous | 0.6 | 0.4 | 3.3±0.6 | 0.9±0. | 0.27±0.08 |
| 12 | Desert | 27.9 | 0.4 | 2.9±0.5 | 0.9±0.1 | 0.31±0.06 |
| 13 | Flooded savanna | 1.1 | 0.3 | 2±0.5 | 0.5±0 | 0.25±0.08 |
| 14 | Mangroves | 0.3 | 0.2 | 2.1±0.6 | 0.4±0.1 | 0.19±0.07 |
| | Globe | 132.4 | 47.3 | 566.2±174.3 | 141.6±31.9 | 0.25±0.10 |

Table 4. Mean and median *R:S* from observations and predicted in this study. The mean *R:S* is the arithmetic average of individual R:S across site level observations (Obs) or gridcells (Predicted). The median is the 50$^{th}$ percentile across observations (Obs) or gridcells (Predicted). Note the mean and median *R:S* are different from the weighted *R:S* from the last column of Table 3 which shows the ratio between total root biomass and shoot biomass. The weighted *R:S* is weighted by biomass while the mean and median are not weighted by biomass.

| **Biome number** | **Name** | **Mean (Obs)** | **Median (Obs)** | **Mean (Predicted)** | **Median (Predicted)** |
|---|---|---|---|---|---|
| 1 | Tropical moist | 0.37 | 0.32 | 0.26 | 0.24 |
| 2 | Boreal | 0.45 | 0.32 | 0.27 | 0.26 |
| 3 | Tropical savanna | 0.44 | 0.36 | 0.29 | 0.27 |
| 4 | Temperate broadleaf | 0.58 | 0.38 | 0.28 | 0.26 |
| 5 | Temperate coniferous | 0.29 | 0.25 | 0.29 | 0.26 |
| 6 | Tropical dry | | | 0.33 | 0.30 |
| 7 | Tundra | | | 0.34 | 0.29 |
| 8 | Temperate savanna | 0.74 | 0.45 | 0.36 | 0.33 |
| 9 | Montane | 0.42 | 0.42 | 0.41 | 0.35 |
| 10 | Mediterranean | 0.43 | 0.35 | 0.39 | 0.35 |
| 11 | Tropical coniferous | 0.671 | 0.55 | 0.35 | 0.31 |
| 12 | Desert | | | 0.40 | 0.35 |
| 13 | Flooded savanna | | | 0.33 | 0.32 |
| 14 | Mangroves | 0.47 | 0.40 | 0.26 | 0.25 |
| | Globe | 0.50 | 0.36 | 0.29 | 0.26 |

### Comparison with published results

There are few studies quantifying large scale vegetation root biomass. We searched through literature and compared our study with earlier studies[8-11]. We grouped here forests into mega-biomes of tropical, temperate and boreal systems to enable a comparison between different studies that used different forest biome definitions and areas (see Table 5). The three mega-biomes together take a share of ~68% of the global total root biomass[8] (forest and non-forest together), and they are also more commonly reported and convenient to compare across studies.

It is unclear whether forest in tropical/subtropical grasslands, savannas and shrublands (Biome 3, Figure 2) should be treated as a tropical forest across studies, forest in temperate grasslands/savannas and shrublands (Biome 8) treated as a temperate forest, and forest in tundra (Biome 7) as a boreal forest. We therefore conducted two series of comparisons with and without the above-mentioned ambiguous forest classes. In series 1 (S1), Biomes 1, 6, 11 and 3 (Biome distribution is displayed in Figure 2) are aggregated to represent tropical systems; Biomes 3, 5, 8 are grouped into temperate forest; and Biomes 6 and 7 are grouped into boreal forest. In series 2 (S2), we grouped Biomes 1,2,3 into tropical forest, Biomes 4 and 5 into temperate forest and Biomes 6 as boreal forest. Together, root biomass from tropical, temperate and boreal forests is 44-183% higher in earlier studies than in S1 and 65-226% higher than in S2 (Table 5).

This over-estimation from earlier studies is largely explained by their over-estimation of shoot biomass. To demonstrate this, we compiled additional studies (Table 6) that reported shoot biomass at the global, tropical, temperate and boreal forests.

The global forest root biomass ranges between 154 – 210 Pg if root biomass was upscaled through different allometric equations collected from literature (Table 7). A prediction of root biomass after fitting our site-level data with an allometric equation (fitted equation: $R = 0.289S^{0.974}$, $R^2 = 0.79$, Table 7) yielded a global forest root biomass of 155 Pg (tree-level-upscaling) or 172 Pg (stand-level-upscaling), which is larger than 147 Pg from the RF up-scaling model. By stand-level-upscaling, we followed the practice in literature[12,13] and assumed an allometric equation is equally applicable to stand level data (weight per area) despite being derived from individual-level data. Root biomass density (weight per area) was directly estimated from GlobBiomass-AGB[7] shoot biomass density through the allometric equations. In tree-level upscaling, similarly to the RF upscaling procedure, GlobBiomass-AGB[7] shoot biomass density was firstly downscaled to individual tree level through tree density[14]. Allometric equations were applied to estimate tree level root biomass (weight per plant), which is then transferred into per area level through the same tree density. Whether it is upscaled from the individual-tree-level or the stand-level is unlikely to explain the overestimation as there is no systematic difference between these two approaches (Table 7).

Table 5. Comparison between studies quantifying root biomass in tropical, temperate and boreal forests. This table adds upon Table 1 in the main text with shoot biomass, land area, biomass density and *R:S*.

| | | This study[S1] | This study[S2] | Jackson1997[8] | Saugier2001[15] | Robinson2007[11] |
|---|---|---|---|---|---|---|
| Method | | Machine learning | Machine learning | Biome average root biomass density, area | Biome average *R:S* ratio, shoot biomass density, area | Biome average *R:S* ratio, shoot biomass density, area |
| Root biomass | Tropical (Tr, Pg) | 92 | 76 | 114 | 147 | 246 |
| | Temperate (Te, Pg) | 26 | 25 | 51 | 59 | 98 |
| | Boreal (Bo, Pg) | 21 | 20 | 35 | 30 | 50 |
| | Tr + Te + Bo (Pg) | 139 | 121 | 200 | 236 | 394 |
| | $RD_{S1}$* | 0% | | 44% | 70% | 183% |
| | $RD_{S2}$& | | 0% | 65% | 95% | 226% |
| Shoot biomass (Pg) | Tropical | 364 | 312 | | 532 | 532 |
| | Temperate | 102.9 | 98.2 | | 218.4 | 218.4 |
| | Boreal | 81.4 | 77.5 | | 83.6 | 83.6 |
| Forest area ($10^6$ $km^2$) | Tropical | 24.1 | 17.4 | 24.5 | 17.5 | 17.5 |
| | Temperate | 9 | 8.3 | 12 | 10.4 | 10.4 |
| | Boreal | 12.1 | 11.2 | 12 | 13.7 | 11.2 |
| Root density (kg/m2) | Tropical | 3.8 | 4.4 | 4.6 | 8.4 | 14.0 |
| | Temperate | 2.9 | 3.0 | 4.2 | 5.7 | 9.4 |
| | Boreal | 1.7 | 1.8 | 2.9 | 2.2 | 4.5 |
| Shoot density (kg/m2) | Tropical | 15.1 | 17.9 | | 30.4 | 30.4 |
| | Temperate | 11.4 | 11.8 | | 21 | 21 |
| | Boreal | 6.73 | 6.9 | | 6.1 | 7.5 |
| Average *R:S* | Tropical | 0.25 | 0.24 | | 0.28 | 0.46 |
| | Temperate | 0.25 | 0.25 | | 0.26 | 0.45 |
| | Boreal | 0.26 | 0.26 | | 0.37 | 0.6 |

S1. Tropical moist forest (Biome 1), tropical dry forest (Biome 6), tropical/subtropical coniferous forest (Biome 11) and forest in tropical/subtropical grasslands/savannas and shrublands (Biome 3) are aggregated to represent tropical systems (Tr). Temperate broadleaf/mixed forest (Biome 4), temperate coniferous forest (Biome 5) and forest in temperate grasslands/savannas and shrublands (Biome 8) are merged together as temperate systems (Te). Boreal forest (Biome 2) and woody plants in tundra region (Biome 7) are aggregated as boreal forest (Bo). Biome classification is from The Nature Conservancy[1] and is shown in Figure 2.
S2. Tropical systems (Tr): Biomes 1,6,11; Temperate systems (Te) : Biomes 4,5; Boreal systems (Bo) : Biome 2.
* $RD_{S1}$, the relative difference of Tr + Te + Bo between this study (S1) and previous quantifications. $RD_{S1}$ = (previous study – this study)/this study x 100%. For example, in the column with the head Jackson, $RD_{S1}$ = (200-139)/139*100% = 44%.
& $RD_{S2}$, the same as $RD_{S1}$, but with the S2 definition of tropical, temperate and boreal systems.

Table 6. Comparison between shoot biomass used in this study[7] and other estimates for tropical, temperate, boreal forests and the globe.

| | | This study[S1] | This study[S2] | Pan2011[16,17] | Saatchi[12] | Liu2015[3] | Bacchini2017[18] | Hu2016[2] |
|---|---|---|---|---|---|---|---|---|
| Method | | GlobBiomass-AGB | GlobBiomass-AGB | Inventory | Satellite | Satellite VOD | Satellite | Satellite LiDAR |
| Time | | Current | Current | Current | ~2000 | ~2000 | ~2007/8 | Current |
| Shoot biomass (Pg) | Tropical | 364 | 312 | 410 | 346-424 | 360-416 | 318 | |
| | Temperate | 102.9 | 98.2 | 88 | | 74-132 | | |
| | Boreal | 81.4 | 77.5 | 72.4 | | 48-78 | | |
| | Globe | 566 | 566 | | | | | 533 |

S1. Tropical moist forest (Biome 1), tropical dry forest (Biome 6), tropical/subtropical coniferous forest (Biome 11) and forest in tropical/subtropical grasslands/savannas and shrublands (Biome 3) are aggregated to represent tropical systems (Tr). Temperate broadleaf/mixed forest (Biome 4), temperate coniferous forest (Biome 5) and forest in temperate grasslands/savannas and shrublands (Biome 8) are merged together as temperate systems (Te). Boreal forest (Biome 2) and woody plants in tundra region (Biome 7) are aggregated as boreal forest (Bo). Biome classification is from The Nature Conservancy[1] and is shown in Figure 2.
S2. Tropical systems (Tr): Biomes 1,6,11; Temperate systems (Te) : Biomes 4,5; Boreal systems (Bo) : Biome 2.

Table 7. Global forest root biomass estimated from allometric equations.

| | **Fit** | **Jiang[19]** | **Niklas[20]** | **Robinson[10]** | **Cairns[21]** |
|---|---|---|---|---|---|
| α | 0.289 | 0.332 | 0.372 | 0.384 | 0.338 |
| β | 0.974 | 0.920 | 0.924 | 0.954 | 0.926 |
| Global Total[t] (Pg) | 155 | 165 | 186 | 199 | 167 |
| Global Total[s] (Pg) | 172 | 154 | 176 | 210 | 161 |

Fit: Observed root (*R*) and shoot (*S*) biomass were fitted into an allometric equation, $R = \alpha S^{\beta}$ where $\alpha$ and $\beta$ are allometric coefficients.
Jiang, Niklas and Robinson: coefficients of the allometric equation were taken from corresponding literature.
[t]: tree-based estimation. GlobBiomass-AGB shoot biomass was firstly transferred to individual tree level through tree density. Tree level root biomass was estimated from the allometric equation and the derived tree level shoot biomass. Tree level root biomass was then transferred into per area level through tree density. This approach takes the similar procedure as the machine learning approach.
[s]: stand-based estimation. Per area root biomass was directly estimated from GlobBiomass-AGB shoot biomass through the allometric equation. This approach mimics practice in literature[12,13].

### Preliminary estimation of fine root biomass

Broadly speaking, leave and fine root biomass are highly linked[22]. Ref[22] derived an relationship between annual leaf biomass production and annual root biomass production (Table 1 of Ref[22]). Assuming an annual turnover of leaves and fine root, we approximate fine root biomass through above mentioned relationship and leaf biomass. Leaf biomass is estimated through the remote sensed leaf area index (LAI)[23,24] and the observation-based leaf mass per area (or the inverse of specific leaf area)[25]. We apply two LAI dataset, the GIMMS3g[24] and the GlobMAP[23]. We estimate total global fine root in forest (with 15% canopy cover threshold as in the main text) to be 6.7 Pg (GIMMS3g) or 7.7 Pg (GlobMAP). We acknowledge leaf and fine root may not be in sync [26] temporally and/or locally. Our estimation here is preliminary that worth improvement with better understanding of fine root in the future.

### Arithmetic mean *R:S* is always larger than shoot-biomass weighted mean *R:S*

The general form of the allometric equation is given by:

$$R/S = \alpha S^{\beta - 1} \qquad \text{(SI1)}$$

We prove here that if root and shoot biomass are related by Equation SI1, the arithmetic mean *R:S* is always larger than the biomass weighted mean. Suppose that we have two classes of trees or forest stands that differ in shoot biomass, one with size *x*, and the other is *y*. We assume the number of *x* is *m* if we look at the individual-tree-level, or the area is *m* if we look at the stand or larger level, and *n* is the number or area of *y*.

The (shoot) biomass weighted mean *R:S* is:

$$\frac{\alpha m x^{\beta} + \alpha n y^{\beta}}{mx + ny}$$

The arithmetic mean *R:S* is:

$$\frac{\alpha m x^{\beta-1} + \alpha n y^{\beta-1}}{m + n}$$

The difference between the weighted and arithmetic mean is:

$$deltaMean = \frac{\alpha m x^{\beta} + \alpha n y^{\beta}}{mx + ny} - \frac{\alpha m x^{\beta-1} + \alpha n y^{\beta-1}}{m + n}$$

By algebraic transformations, this equation can be transformed into:

$$deltaMean = \frac{\alpha m n}{(m+n)(mx+ny)}(x-y)\left(x^{\beta-1} - y^{\beta-1}\right) \qquad (SI2)$$

Since we have $\alpha, m, n, x, y > 0,$ Equation SI2 tells $if\ \beta = 1, deltaMean = 0; if\ \beta < 1, deltaMean < 0;\ if\ \beta > 1, deltaMean > 0.$ Both theory and empirical evidence across world's forests lead to *R:S* vs. S relationships like Equation SI1 with $\beta < 1$,[9,27,28], which proves that the arithmetic mean *R:S* always overestimate the (shoot) biomass weighted mean *R:S*.

**Allometric upscaling overestimates *R:S* at 1km resolution**

If we assume root and shoot biomass follow a universal allometric equation at different scales (Equation SI1), we show here we would always overestimate root biomass from the average shoot biomass at the pixel level. Here, we take the 1-km resolution as an example and upscaling to other resolutions follow the same logic. We start from upscaling from individual trees and discuss later the case for the stand-level. Suppose we have two classes of trees or forest stands that differ in shoot biomass, one with size *x*, and the other is $y$. In tropical forest, the number of individuals ($N$) generally follows a tight power law distribution, with the dominant power function of the form $d^{-(\theta+1)}$ , where $d$ is the tree diameter and $\theta$ is related to the allometric exponent of the crown area to diameter[29], which is relatively consistent across tropical forests. Reported value of $\theta$ is around 1.27-1.31. In temperate or boreal forests, sometimes there may lack the above power law size structure, and we will discuss this case later. The relationship between tree diameter and biomass is highly conserved, with idealized trees exhibiting a general allometric function where $AGB \propto d^{\omega}$ [30]. The range of $\omega$ is between 1.1 and 3.37 from China's tree biomass equation database which consists of 5,924 biomass component equations for nearly 200 species. Together,

$$N = \mu \, AGB^{-\frac{\theta+1}{\omega}}$$

where $\mu$ is a parameter with a positive value. We use $\gamma$ to replace $\frac{\theta+1}{\omega}$ for simplicity, and can write

$$N = \mu \, AGB^{-\gamma}$$

The real *R:S* ratio is,

$$RS_{real} = \frac{\alpha\mu x^{\beta-\gamma} + \alpha\mu y^{\beta-\gamma}}{\mu x^{1-\gamma} + \mu y^{1-\gamma}}$$

Which is the same as:

$$RS_{real} = \frac{\alpha(x^{\beta-\gamma} + y^{\beta-\gamma})}{x^{1-\gamma} + y^{1-\gamma}}$$

The estimated *R:S* is:

$$RS_{esti} = \alpha(\frac{\mu x^{1-\gamma} + \mu y^{1-\gamma}}{\mu x^{-\gamma} + \mu y^{-\gamma}})^{\beta-1}$$

Which is the same as:

$$RS_{esti} = \alpha(\frac{x^{1-\gamma} + y^{1-\gamma}}{x^{-\gamma} + y^{-\gamma}})^{\beta-1}$$

Therefore, the difference between estimated and real *R:S* is,

$$deltaRS = RS_{esti} - RS_{real} = \alpha\left(\frac{\mu x^{1-\gamma} + \mu y^{1-\gamma}}{\mu x^{-\gamma} + \mu y^{-\gamma}}\right)^{\beta-1} - \frac{\alpha\left(x^{\beta-\gamma} + y^{\beta-\gamma}\right)}{x^{1-\gamma} + y^{1-\gamma}} \qquad (SI3)$$

With the condition $\beta < 1, \alpha > 0, \mu > 0, \; x > 0, y > 0, \gamma > 0,$ $deltaRS$ is always bigger than 0, as shown in Figures 10, 11 numerically.

For forests without the power law structure or when we upscale from the stand-level measurement, we use *m* and *n* to denote the number of trees or the area of stands with the size of shoot biomass *x* and *y*.

The difference between estimated and real *R:S* is,

$$deltaRS = RS_{esti} - RS_{real} = \alpha\left(\frac{mx + ny}{m + n}\right)^{\beta-1} - \frac{\alpha\left(mx^{\beta} + ny^{\beta}\right)}{mx + ny} \qquad (SI4)$$

With the condition $\beta < 1, \alpha > 0, \mu > 0, \; x > 0, y > 0, m > 0, n > 0, \gamma > 0,$ $deltaRS$ is always bigger than 0 as illustrated in Figures 12, 13 numerically.

The magnitude of overestimation is related to $\beta, \alpha, \mu, x, y, m, n$ (or $\gamma$ in case of forests with power law size structure).

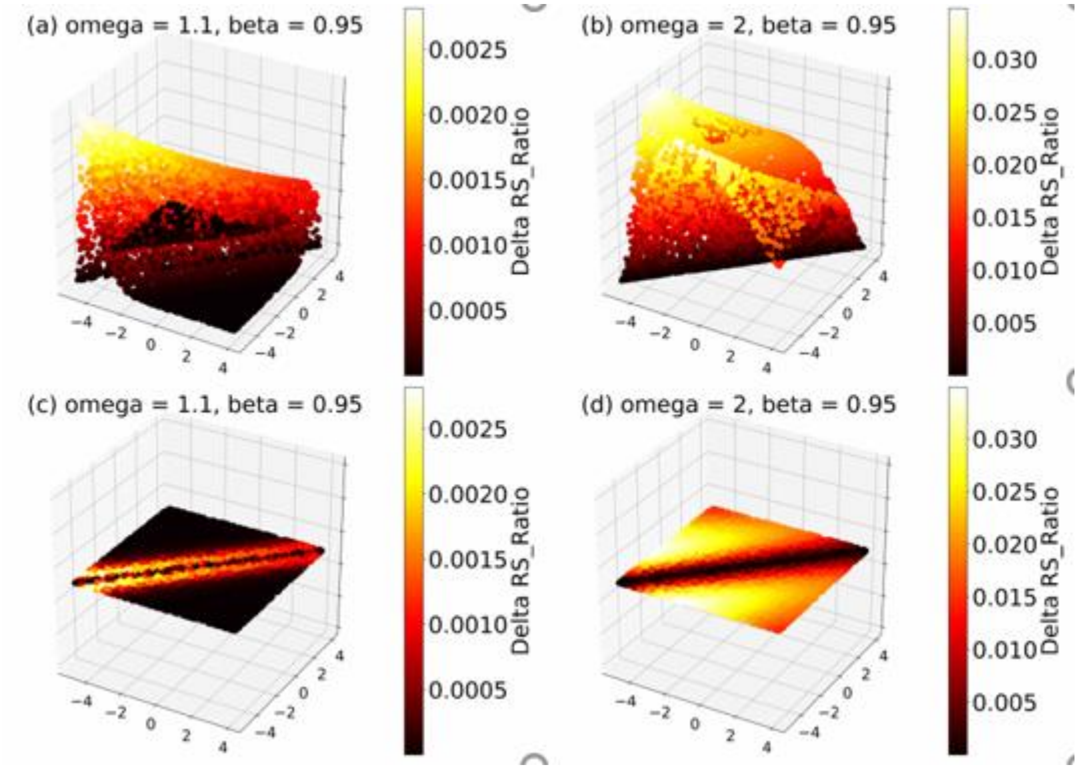


Figure 10, $deltaRS$ in responses to changes in tree sizes in *x* (*x*-axis) and *y* (*y*-axis). Size *x* and size *y* are randomly chosen with $\log x, logy \in [-5,4]$. Here we fix $\alpha$ *and* $\theta$ with typical values $\alpha = 0.31$, $\theta = 1.3$. (a) and (c) show $deltaRS$ with ω=1.1, β=0.95. (b) and (d) show $deltaRS$ with ω=2, β=0.95. (a) and (b) display $deltaRS$ in a 3-dimentional space and the (c) and (d) are corresponding projections into the *x*-*y* space. $deltaRS$ is always bigger than 0 with different values of $x, y, \alpha, \theta, \omega, \beta$ in literature. We choose fixed values for demonstration purpose here. See Equation SI3 for details.

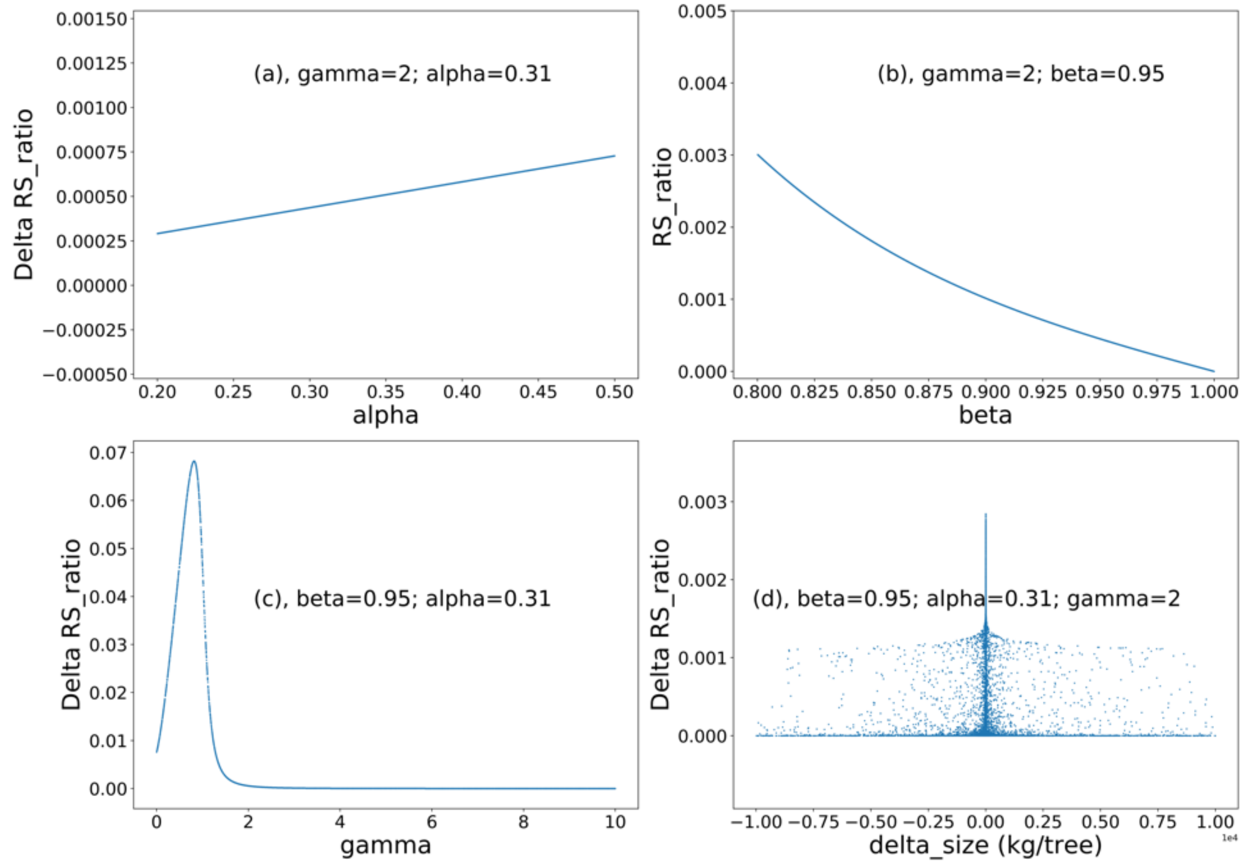


Figure 11. $deltaRS$ in responses to changes in α (a, alpha), β (b, beta), γ (c, gamma) and difference in tree size (d, delta_size). In panels (a), (b) and (c), the parameter in *x-axis* varies in a range that is broader than typically reported in literature while other parameters are fixed at a typical value. Panel (d) shows changes in $deltaRS$ in response to differences in size *x* and size *y* where size *x* and size *y* are randomly generated with a uniform distribution of *logx* and *logy* with $\log x, logy \in [-5,4]$. Note, in (d) Delta RS_ratio = 0 when delta_size = 0, but varies largely in a small region around 0. See Equation SI3 for details.

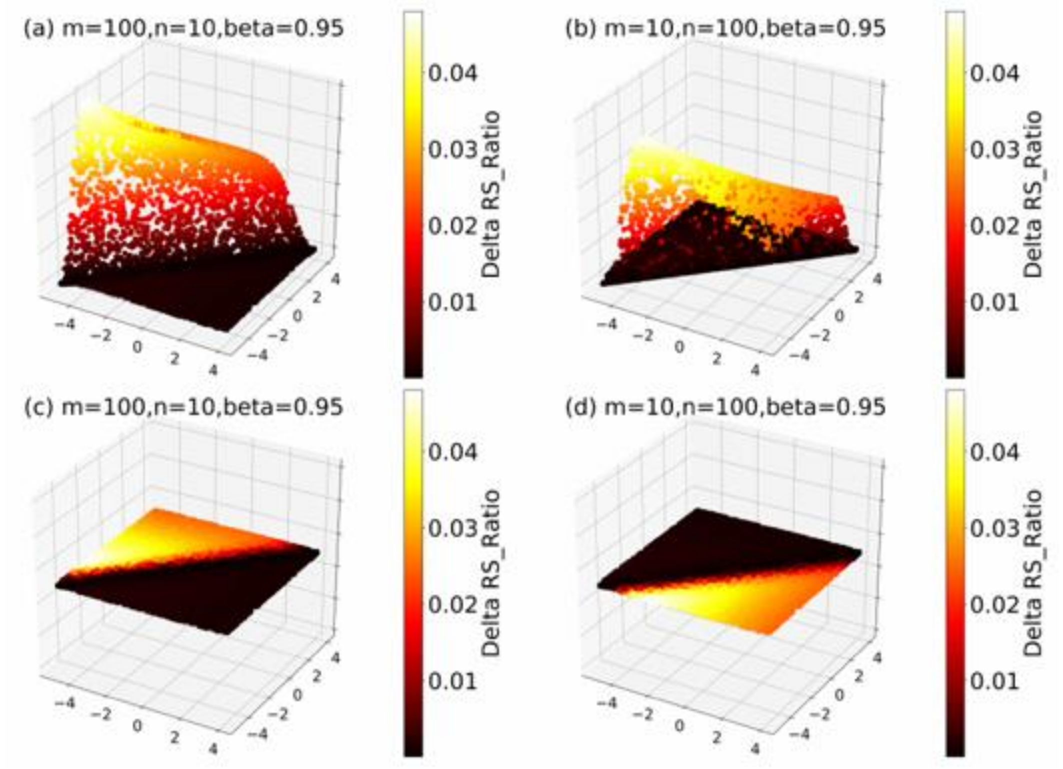


Figure 12. $deltaRS$ in responses to changes in tree sizes in *x* (*x*-axis) and *y* (*y*-axis). Size *x* and size *y* are randomly chosen with $\log x, logy \in [-5,4]$. Here we fix $\alpha$ $and$ $\theta$ with typical values $\alpha = 0.31,\ \theta = 1.3.$ (a) and (c) show $deltaRS$ with *m*=100, *n*=10, β=0.95. (b) and (d) show $deltaRS$ with *m*=10,*n*=100, β=0.95. (a) and (b) display $deltaRS$ in a 3-dimentional space and (c) and (d) are their corresponding projection into the *x*-*y* space. $deltaRS$ is always bigger than 0 with different values of $x, y, \alpha, \theta, m, n, \beta$ in literature. We choose fixed values for demonstration purpose here. See Equation SI4 for details.

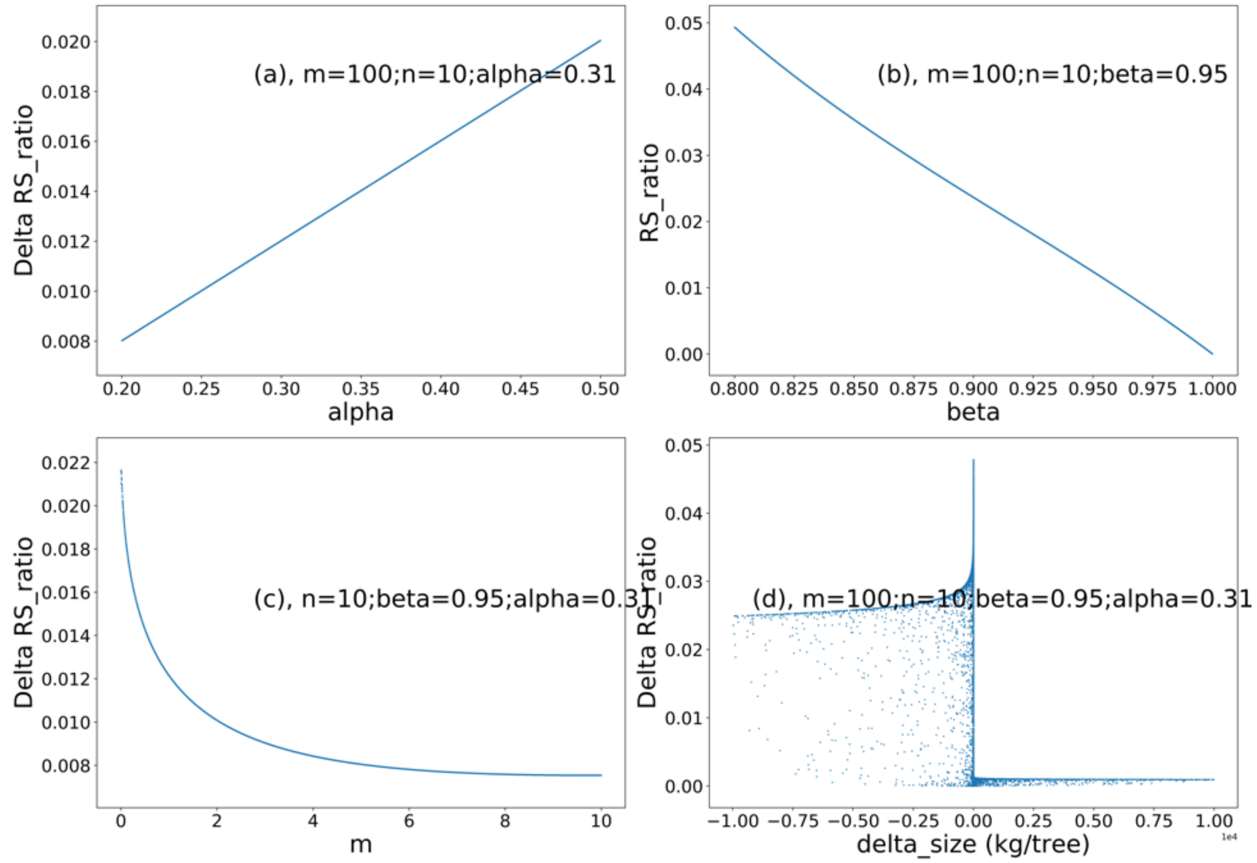


Figure 13. $deltaRS$ in responses to changes in α (a, alpha), β (b, beta), number of trees or stand area of shoot biomass class *x* (c, *m*) and difference in tree size (d, delta_size). This figure is the same as Figure 10 except the exponent controlling the number of trees (γ) is replaced by the number of trees or stand area of each biomass size (*m* and *n*). Note, in (d) Delta RS_ratio = 0 when delta_size = 0, but varies largely in a small region around 0. See Equation SI4 for details.

**Root biomass prediction with age as a predictor**

When age is fixed as a predictor in the random forest model, the "best" trained model incorporates 14 additional predictors which are shoot biomass, height, soil nitrogen, pH, bulk density, clay content, sand content, base saturation, cation exchange capacity, vapor pressure, mean annual precipitation, mean annual temperature, aridity and water table depth. This model slightly reduced the mean absolute error (MAE = 2.16 vs. 2.18). Global total root biomass from this model is similar to the model without age. The age map is merged from several different sources (see Method), which likely introduce additional uncertainty in our estimation. We therefore prefer the prediction without age as a predictor.
